# Systematic hormone-metabolite network provides insights of high salinity tolerance in *Pongamia pinnata* (L.) pierre

**DOI:** 10.1101/2020.04.28.066050

**Authors:** Sureshbabu Marriboina, Kapil Sharma, Debashree Sengupta, Anurupa Devi Yadavalli, Rameshwar Prasad Sharma, Attipalli Ramachandra Reddy

**Affiliations:** Department of Plant Sciences, School of Life Sciences, University of Hyderabad, Hyderabad-500046, India; Repository of Tomato Genomics Resources, Department of Plant Sciences, School of Life Sciences, University of Hyderabad, Hyderabad-500046, India; Department of Animal Biology, School of Life Sciences, University of Hyderabad, Hyderabad 500046, India

**Author notes:** **Corresponding author:** Prof. A. R. Reddy, Department of Plant Sciences, School of Life Sciences, University of Hyderabad, Hyderabad-500046, Telangana, India.

**Keywords:** Carbon exchange rate, correlation studies, hierarchical cluster analysis, hormone-metabolite correlation, metabolomics, metabolite-metabolite correlation, Na^+^-localization, phytohormone, transporter genes

## Abstract

Salinity stress results significant losses in plant productivity, and loss of cultivable lands. Although *Pongamia pinnata* is reported to be a salt tolerant semiarid tree crop, the adaptive mechanisms to saline environment are elusive. The present investigation describes alterations in hormonal and metabolic responses in correlation with physiological and molecular variations in leaves and roots of Pongamia at sea salinity level (3% NaCl) for 8 days. At physiological level, salinity induced adjustments in plant morphology, leaf gas exchange and ion accumulation patterns were observed. Our study also revealed that phytohormones including JAs and ABA play crucial role in promoting the salt adaptive strategies such as apoplasmic Na^+^ sequestration and cell wall lignification in leaves and roots of Pongamia. Correlation studies demonstrated that hormones including ABA, JAs and SA showed a positive interaction with selective compatible metabolites (sugars, polyols and organic acids) to aid in maintaining osmotic balance and conferring salt tolerance to Pongamia. At the molecular level, our data showed that differential expression of transporter genes as well as antioxidant genes regulate the ionic and ROS homeostasis in Pongamia. Collectively, these results shed new insights on an integrated physiological, structural, molecular and metabolic adaptations conferring salinity tolerance to Pongamia.

**High light:** Our data, for the first time, provide new insights for an integrated molecular and metabolic adaptation conferring salinity tolerance in Pongamia. The present investigation describes alterations in hormonal and metabolic responses in correlation with physiological and molecular variations in Pongamia at sea salinity level (3% NaCl) for 8 days.

## Introduction

Soil salinization is a major environmental constraint to limit plant growth and production which is closely associated with arable land degradation (Shahid et al., 2018; Marriboina Attipalli, 2020a). Approximately 1.5 million hectares of cultivable land are becoming saline marginal lands by every year because of high salinity levels and nearly 50% of arable lands will be lost by year 2050 (Hossain, 2019). Global climate change, increasing population and excessive irrigation are further limiting the availability of cultivable land for crop production (Raza et al., 2019). To date, attempts are being made to extend the crop productivity on saline lands. However, progress of these attempts is greatly hampered by the genetic complexity of salt tolerance, which largely depends on physiological and genetic diversity of the plant and spatio-temporal heterogeneity of soil salinity (Morton et al., 2019). To address this issue, several plant species were introduced to rehabilitate the saline lands and certain economically important nitrogen fixing biofuel tree species are of immense importance not only for sustenance to saline marginal lands but also economic gain towards saline lands (Samuel et al., 2013; Hanin et al., 2016; Marriboina Attipalli, 2020a).

A comprehensive understanding of physiological, hormonal and molecular adaptive mechanisms is crucial to cultivate these tree species on saline marginal lands (Quinn et al., 2015). Plants growing in saline soils prevent the excess Na^+^ ion disposition in the leaves in order to protect the photosynthetic machinery from salt-induced damage. The decrease in net CO_2_ assimilation rate and optimum quantum yield of PSII (Fv/Fm) might substantiate the leaf performance under salt induced drought stress. Calcium ion (Ca^2+^) is known as an intracellular second messenger and plays an important role in plant growth and development. It also plays an essential role in amelioration of sodium toxicity through activating several Ca^2+^ responsive genes and channels (Thor, 2019). In response to salt stress, plant produces several phytohormones such as ABA, JA, MeJA, zeatin, IAA, IBA and SA, which plays crucial role in sustaining its growth under extreme saline conditions. ABA is well-known stress induced phytohormone, critical for plants growth and regulating numerous downstream signalling responses (Tuteja, 2007). ABA causes stomatal closure to prevent excess water evaporation and regulate root growth under salinity stress (Zelm et al., 2020). Auxins and cytokinins are growth promoting phytohormones interacts to regulate various growth and developmental process such as cell division, elongation and differentiation. Salt induced endogenous accumulation of cytokinin improves the salt tolerance in crop species by delaying leaf senescence and marinating photosynthetic capacity (Liu et al., 2012; Gloan et al., 2017). Upon salt stress, the raise in endogenous SA levels can cause a significant reduction in the ROS and Na^+^ accumulation across the plant, whereas SA deficient plant produced an elevated levels of superoxide and H_2_O_2_ (Yang et al., 2004). According to Sahoo et al., (2014), the perfect harmony among phytohormones played a significant role in improving the salt tolerance in rice. However, the synergistic and antagonistic interactions between phytohormones are mostly depending on plant species and type of stress imposed on plants, but their interactions are still not clearly understood (Gupta et al., 2017). To combat against salinity induced ROS damage, plants adapted jasmonates directed anthocyanin accumulation to mitigate its negative effects (Ali Baek, 2020). Further, JAs can positively regulate the endogenous ABA level, together regulate the guard cell movement during salt stress (Siddiqi Husen, 2019). JA and ABA together regulates antioxidant status of the cell to enhance the survivability of plant towards osmotic stress. Additionally, JA and SA positively regulate the several protein coding genes which are responsible for plant salt tolerance (Wang et al., 2020). Plant pre-treated with SA alleviates salinity stress by decreasing Na^+^ transport and by increasing H^+^-ATPase activity (Jayakannan et al., 2013; Gharb et al., 2018). Excessive deposition of salts in the cell and celluar compartments causes membrane depolarization. Counteract to the salt-induced membrane depolarization ABA regulates expression of numerous vacuolar and plasma membrane transporters such as vacuolar H^+^-inorganic pyrophosphatase, vacuolar H^+^-ATPase, NHX1, V-PPase, and PM-H^+^-ATPase pumps (Fukuda and Tanka, 2006). According to Shahzad et al., (2015), exogenous application of JAs on maize improved salt tolerance by regulating Na^+^ ion uptake at the root level. Proton pumps and cation channels such as H^+^-ATPase pumps, CHXs and CCXs were involved in maintaining the membrane potential under salt stress conditions (Falhof et al., 2016; Li et al., 2016; Liu et al., 2017). Importantly, plant induces the expression of several isoforms of NHXs namely SOS1, NHX1, NHX2, NHX3 and NHX6 under salt stress to regulate Na^+^ fluxes in and out of the cell (Dragwidge et al., 2018). Upon salt stress, plants activate a complex antioxidant defense mechanism to minimize oxidative stress damage by ROS under salt stress conditions (Xie et al., 2019).

Population and industrialization pressure has increased the demand for land and fossil fuel resources. In addition, there is increasing demand for renewable energy resources due to the fast depletion of fossil fuel resources. For the first time, we report here the mechanisms of salinity tolerance in Pongamia at molecular level with the help of hormonal, metabolomics, gene expression and computational approaches. Further, the assessment of tissue specific-phytohormone profiling elucidates the key role of specific hormones in conferring the tissue-specific associated mechanisms of salt tolerance in Pongamia. In addition, the correlation studies between the phytohormones enable to identify the crosstalk between phytohormones, which may regulating the growth and development in Pongamia under salt stress (Maury et al., 2019). Time-course metabolic profiling and correlations under salt stress in Pongamia would certainly contribute to understand the biochemical changes involving metabolic pathways, which is crucial in plant adaptation to salinity stress conditions. The interaction studies between hormones and metabolites should certainly create new opportunities for the discovery of hormone-metabolite associates, which are very crucial to understand stress tolerant mechanisms (Cao et al., 2017). The present study provides an evidence for the hormone-metabolite interactions as well as novel hormone-metabolite associated signalling pathways to understand high salinity tolerance mechanisms in *Pongamia pinnata*.

## Materials and methods

### Plant material, growth conditions and NaCl treatment

Pongamia seeds (accession TOIL 12) were obtained from Tree India Limited (TOIL), Zaheerabad, Hyderabad, Telangana. Freshly collected seeds were sterilized with 1% (v/v) hypochlorite solution for 5 min. Further, the seeds were kept for germination on moist cotton bed at 25°C in dark for 10 days after thoroughly washed with autoclaved distilled water. Germinated seeds were grown hydroponically in Hoagland No. 2 basal salt mixture (Himedia) solution (pH 5.75 ± 0.02) for 30 days. The seeds were kept for germination on moist cotton bed for 10 days at 25°C in dark after thoroughly washed with autoclaved distilled water. After radicle emergence, the germinated seeds were placed just above the nutrient medium (Hoagland No. 2 basal salt mixture (Himedia) solution (pH 5.75 ± 0.02)) level with help of parafilm in 50 ml falcon tubes (Genaxy) for 10 days. Further, the plants were grown initially in long cylindrical glass tubes (5 cm diameter X 30 cm length) for 10 days and then transferred to long cylindrical glass tubes (5 cm diameter X 60 cm length) before giving the salt stress treatment. The glass apparatus were designed in such a way to grow the plants without root limitation. We maintained following culture conditions 24°C temperature, 16 h light and 8 h dark, and relative humidity maintained approximately at 60%. In order to avoid hypoxic conditions nutrient solution was renewed on daily basis. Salinity stress treatment was given according to Marriboina Attipalli, (2020b). For salinity stress, two different salt concentrations (300 mM NaCl and 500 mM NaCl (sea water equivalents)) were used. For stress treatment, 30 days old plants (n = 20 to 30) were chosen and exposed to 300 and 500 mM NaCl stress with increment of 100 mM NaCl per day (Marriboina Attipalli, 2020b). Control plants were maintained in a fresh Hoagland’s solution. Plants were harvested at an interval of 1, 4 and 8 days. Fresh weights of leaves and roots were measured immediately after harvest. Dry weights of control and treated plant tissues were determined after drying at 70°C for 3 days. Control and treated samples were harvested at each time intervals, flash frozen in liquid nitrogen and stored at −80°C till analysis.

### Leaf gas exchange and chlorophyll fluorescence parameters

Leaf net photosynthetic capacity was measured with the flow-through Q-box CO650 (Qubit Systems). All measurements were performed on fully expended 2^nd^ and 3^rd^ leaf of the plant between 9:00 to 11:00 h under the following conditions: a saturating photosynthetic active radiation (PAR) of 1600 *µ*mol m^-2^. s^-1^ supplied by LED light source (A113), attached to leaf chamber continuously flushed with 350 ppm CO_2_ containing air at a flow rate of 50 ml. min^-1^, air temperature of 25-26°C and relative humidity of 55-60%. The measurements were taken after CO_2_ output stabilized, and CO_2_ exchange rate was calculated with the Logger Pro software (Qubit Systems). The details of the maximum quantum yield (Fv/Fm) and Chl*a* fluorescence analysis was performed by using a portable mini-PAM chlorophyll fluorometer (Walz, Germany) fitted with standard fiber optic and leaf clip holder. To measure Chl*a* fluorescence, leaves were kept in dark incubation for 30 min. For analysis, we used the following conditions: intensities of measuring light (<0.1 *µ*mol photons m^-2^. S^-1^), saturating pulse (3000 *µ*mol photons m^-2^. S^-1^), actinic light (170 *µ*mol photons m^-2^. S^-1^) and far-red light (7 *µ*mol photons m^-2^. S^-1^) respectively. The duration of actinic light and saturation pulse were 0.8 and 30 s. Chlorophyll *a* and *b* were estimated using Arnon, (1949) method.

### Quantification of Na^+^, K^+^, Ca^2+^ and Cl^-^ ions

Ion content in leaves and roots of control and salt treated plants were determined as described in Munns et al., (2010) with some minor modifications. The collected samples were oven dried at 70°C for 3 days and ground finely with motor and pestle. For ion extraction, 1 g of finely ground sample powder was taken in 5 ml of aqua regia (concentrated HCl: HNO_3_, 3:1 v/v) and boiled at 95°C for 60 min. The resulting solution was analysed by using an atomic absorption spectrophotometer (GBC 932, Braeside, Australia). Chloride ion concentration was determined by titrimetric method according to Mohr’s method (Korkmaz, 2001).

### Visualization of Na^+^ ions through confocal laser scanning microscopy (CLSM)

Freshly collected leaves and roots were segmented into 1 cm sections and incubated with 2.5% glutaraldehyde solution in 0.1 M MOPS buffer overnight at 4°C. For sodium illumination, tissues were stained with 5 *µ*M Na^+^ specific probe CoroNa-Green AM (Invitrogen) in the presence of 0.02% pluronic acid in 50 mM MOPS (pH 7.0) for overnight at room temperature. Further, the segments of leaves and roots were thoroughly washed with 50 mM MOPS (pH 7.0) several times. Thin sections were performed with razor blade and quickly immersed in cell wall stain propidium iodide (Invitrogen) for 15 min. The sections were examined under CSLM (Leica TCS SP2 with AOBS, Heidelberg, GmbH, Germany). All measurements were performed as described by Marriboina Attipalli, (2020b) with some minor modifications. The cytosolic and vacuolar Na^+^ fluorescence were calculated by LCS software (Heidelberg, GmbH, Germany).

### LC-MS analysis

The endogenous levels of phytohormones were quantified by LC-MS using a protocol from Pan et al., (2010) with some modifications. Freshly harvested leaves and roots of control and salt-treated plants were finely ground to powder with liquid nitrogen. Approximately 100 mg of powdered tissue was suspended in 500 *µ*l of extraction solvent (2-Propanol:H_2_O:HCl, 2:1:0.002, v/v/v) and kept in thermomixer at 4°C at 500 rpm for 30 min. The above step was repeated with addition of 1 ml ice cold dichloromethane. The resulting mixture was centrifuged at 4°C 13000g for 5 min and collected in fresh centrifuge tube. The sample tubes were placed in speed vac to evaporate extra solvent for 45 min. The final residue was dissolved in 70 *µ*l of ice cold methanol followed by centrifugation at 13000g for 5 min. Finally, samples were taken into a transfer vial and were analysed by using Exactive™Plus Orbitrap mass spectrometer (Thermo Fisher, USA) coupled with UPLC (Waters, Milford, MA, USA).

LC-MS analysis was performed on an Aquity UPLC™ System equipped with quaternary pump, and auto-sampler to perform hormone analysis. For analysis, we used following conditions: capillary temperature 350°C, sheath gas flow (N2) 35 (arbitrary units), AUX gas flow rate (N2) 10 (arbitrary units), collision gas (N2) 4 (arbitrary units) and capillary voltage 4.5 kV under ultrahigh vacuum 4e-10 mbar. Chromatographic separation was carried out in a Hyperreal GOLD C18 (Thermo Scientific) column (2.1×75 mm, 2.7 μM). Formic acid (0.1%, v/v) and acetonitrile with 0.1% formic acid were used as mobile phase A and B, respectively. A gradient elution program was performed using two solvents system, solvent A and B chromatographic run for 9 min at 20°C. Hormones such as ABA, GA, JA and SA were quantified by using all ion fragmentation (AIF) mode (m/z 50-450) with positive heated electrospray ionization (ESI) in negative ion mode. Similarly, hormones zeatin, IAA, indolebutyric acid (IBA) and methyljasmonate (MeJA) were measured using Turbo Ion spray source in positive ion mode. All phytohormones were quantified by using standard calibration curves.

### GC-MS analysis

The plant primary metabolites were analysed by GC-MS as described by Roessner et al., (2000). 100 mg of freshly collected leaves and roots were ground finely with liquid nitrogen and extracted with 1.4 ml of 100% methanol containing ribitol as internal standard. The mixture was kept at 70°C for 15 min and then mixed vigorously with 1.4 ml of water. Further, the sample was transferred in glass vial (GL-14 Schott Duran) and centrifuged at 4°C for 15 min at 542 rpm. The upper phase was taken in a fresh centrifuge tube and evaporated by using Speed Vac drying for 45 min. Finally, the samples were kept in −80°C for storage until analysis. For derivatization, the residue was dissolved in 80 *µ*l of methoxyamine hydrochloride with pyridine at 30°C for 90 min, followed by 80 *µ*l of MSTFA (N-methyl-N- (trimethylsilyl) trifluoroacetamine) and 20 *µ*l of FAME mix (Sigma, 1 *µ*g. *µ*l^-1^ in hexane) at 30°C for 30 min.

The derivatized sample was analysed on a system LECO-PEGASUS GCXGC-TOF-MS system (LECO Corporation, USA) equipped with 30 m Rxi-5ms column with 0.25 mm internal diameter and 0.25 μm film thickness (Restek, USA). The ion, interference and source of injection temperatures were maintained at 200°C, 225°C and, 250°C, respectively. Chromatographic separation was carried out under following conditions: isothermal heating at 70°C for 5 min, followed by 5°C min^-1^ oven temperature ramp to 290°C and final heating at 290°C for 5 min. The flow rate of carrier gas (helium gas) was adjusted to 1.5 ml. min^-1^. A volume of 1 *µ*l sample was used for analysis and injected in split less mode, and scan mass range 70 to 600 at 2 scans/ s.

All data sets were generated in the form of NETCDF files from ChromaTOF software 4.50.8.0 chromatography version (LECO Corporation, USA) GC-MS and were further exported to MetAlign 3.0 (www.metalign.nl). The MSClust software was used to adjust signal to noise ratio of ≥2, for baseline correction, noise estimation and identification of mass peak (ion-wise mass alignment). Non-repetitive (<3 samples) mass signals were discarded. The MSClust analysis was performed on the above results for the reduction of data and compound mass extraction. Further, the MSClust files were further exported to NISTMS Search v2.2 software for identifying compounds with the NIST (National Institute of Standard and Technology) Library and G□lm Metabolome Database Library (http://gmd.mpimp-golm.mpg.de/). Metabolite identity was given based on the maximum matching factor (MF) value (>700) and least retention index (RI) value. The intensity of each metabolite value normalized with internal standard ribitol value. Finally, only annotated metabolites were taken into consideration for further analysis. Lists of detected metabolites were given in the supplementary data.

### Semi-quantitative PCR and quantitative RT-PCR analysis

In the present study, a total of 34 genes were analysed for their salinity induced expression profiles. Direct primers were used for 22 genes based on available transcriptome given by Sreeharsha et al., (2016) and Huang et al., (2012), while indirect primers were designed for 12 genes based on the available gene sequences of *Glycine max* from SoyKB (www.soykb.org) database since the transcriptome studies showed that *P. pinnata* is closely related to *G. max* (list of primers given in Supplementary Table 7). All genes were amplified by PCR using *P. pinnata* cDNA. The amplicons obtained based on *G. max* indirect primers as well as *P. pinnata* direct primers were sequenced for confirming the identity of target gene. Quantitative PCR analysis was performed on Eppendorf Realplex Master Cycler (Eppendorf, Germany) using KAPASYBRFAST [Mastermix (2×) Universal; KAPA Biosystems] real-time PCR kit following manufacturer’s instructions. For relative quantification analysis, 0.25 *µ*g of RNA template was used, which was extracted from the pool of six biological replicates of both control and salt treated plants by using Sigma spectrumTM Total RNA kit (Sigma, USA) and cDNA synthesis was performed by using the PrimeScriptTM 1^st^ strand cDNA synthesis kit (TAKARA, Japan). Target genes were amplified with the following cycling programme: 1 cycle at 95°C for 2 min, followed by 40 cycles of 30 s at 95°C, 30 s at 60°C and 20 s at 72°C, followed by the dissociation (melting) curve. The intensity of fluorescence was measured by using the Realplex software (Eppendorf, Germany). The limit and efficiency of each primer pair of both target and reference gene was allowed to measure accurate comparison of gene expression using the 2^-ΔΔC^_T_ method for relative quantification (the Applied Biosystems User Bulletin No. 2 (P/N 4303859)) (Livak Schmittgen, 2001). The 18s rRNA was used as reference gene after confirming its consistency expression under salt stress.

### Correlation and statistical analysis

To measure the linear dependence between the variables, Pearson’s correlation coefficient was calculated using R statistical package. Heat maps were generated based on correlation values of metabolites and hormones. Circos plot represents the relationship between hormones and metabolites, which was generated based on correlation values of individual metabolites and hormones. The statistical significant differences of control and salt-treatments were calculated by using one-way ANOVA at P < 0.001, P < 0.01 and P < 0.05.

## Results

### Salinity-induced alterations in growth morphology, gas exchange, chlorophyll fluorescence and ion accumulation patterns of P. pinnata

In order to assess salt tolerance capacity in *P. pinnata*, 30 days old seedlings were exposed to 300 and 500 mM NaCl concentrations for 1, 4 and 8 days. After 8 days of salt treatment (DAS), leaves of 300 mM NaCl treated plants remained healthy and green as that of controls, while 500 mM NaCl stress exposure caused tip burns at 8DAS (Figure 1A). Reason behind selection of the specific time points for this study was that plants were treated with two different salt concentrations (300 mM NaCl and 500 mM NaCl) and kept under observation each day for induction of salt-induced morphological symptoms including yellowing of leaves, necrosis, wilting and shedding of old leaves. However, we couldn’t observe any morphological changes till 7DAS (Days After Salt-treatment) in 500 mM NaCl treated plants. Further, at 8DAS, 500 mM NaCl treated plants started to show wilting symptoms (which was shown in Supplementary Figure S1) and with increasing treatment time further (After 10DAS) these plants were continued to shed off its leaves (Supplementary Figure S1). Interestingly, at this point of time, 300 mM NaCl treated plants did not display any salt-induced morphological symptoms. With the above observation, we concluded that the 8DAS plants could probably give us possible clues regarding salt tolerance mechanisms. We observed that carbon exchange rate (CER) was changed with salt concentration and treatment time (Supplementary Figure 2A). Upon 300 mM NaCl stress, the CER was not changed significantly at 1 and 4DAS, while a significant reduction of ∼50% was observed at 8DAS. In 500 mM NaCl treated plants, the CER values were progressively decreased with the treatment time. However, the levels of chlorophylls pigments were not changed significantly in both 300 and 500 mM salt treated plants at all-time points (Supplementary Figure 2B). Further, 300 mM NaCl treated plants could maintain a constant Fv/Fm values like control plants at all-time points (Supplementary Figure 2C). Conversely, 500 mM NaCl treated plants showed a little decrease in Fv/Fm values at 4 and 8DAS. Similar trend was observed with relative water content (RWC) in leaves and roots of Pongamia. The LRWC was maintained equal to that of controls in 300 mM NaCl treated plants, whereas in 500 mM NaCl treated plants these values tend to decrease progressively with the treatment time (Supplementary Figure 2D). In contrast, the RRWC values did not change significantly in both 300 and 500 mM NaCl treated plants.

**Fig. 1.**
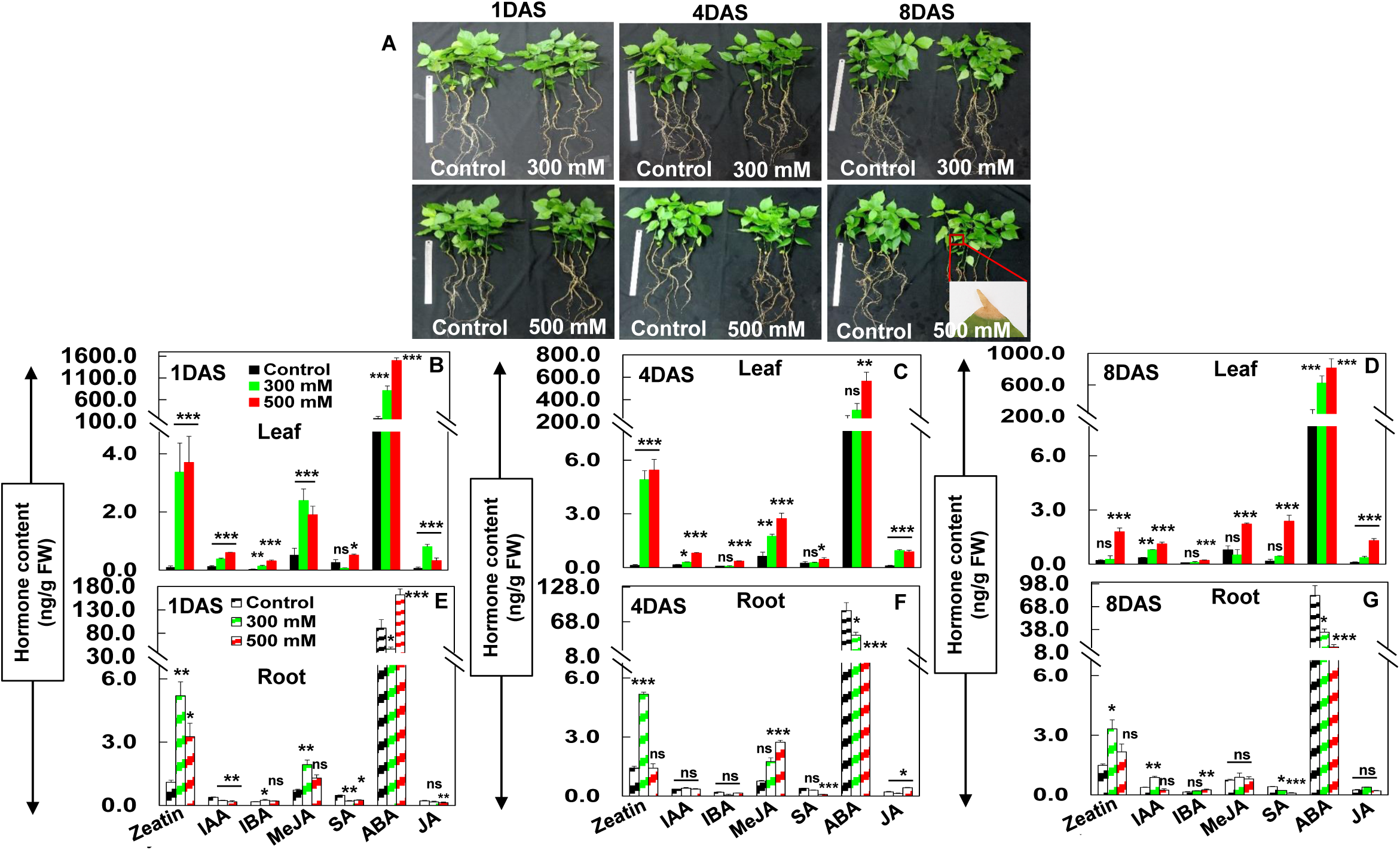
Effect of salt stress on plant morphology and levels of endogenous phytohormones in leaves and roots of *Pongamia pinnata*. (A) Shoot and root morphology of Hoagland’s solution-grown *P. pinnata* plants after 30 days treated with three different salt concentrations 0 (control), 300 and 500 mM NaCl at three different time points 1, 4 and 8DAS. The levels of endogenous hormones (B and I) zeatin, (C and J) IAA, (D and K) IBA, (E and L) MeJA, (F and M) SA, (G and N) ABA and (H and O) JA in leaves and roots of 0 (control), 300 and 500 mM NaCl treated plants at three different time points1, 4 and 8DAS respectively. Error bar represents the mean ± SD (n=6). Two-way ANOVA test was performed to measure P-values ns (not significant), * (P<0.05), ** (P<0.01) and * * * (P<0.001) respectively.

Differences in photosynthetic and morphological responses to salt stress showed in Figure 1A and Supplementary Figure 2A-D might result of difference in their Na^+^ accumulation patterns across the plant, we tested the following hypothesis. As shown in the Supplementary Figure 2E-G, ions such as Na^+^, Cl^-^, and Ca^2+^ content were increased dose dependently in both 300 and 500 mM NaCl treated plants with treatment time. In detail, root showed a significant increase in Na^+^ content when compared to leaves. The peak value of root Na^+^ ion content was about 65 mg. g^-1^ DW in 500 mM NaCl treated plants at 8DAS. Similarly, root Cl^-^ levels were enhanced by ∼2.7, ∼2.7, and ∼1.4-fold in 300 mM NaCl treated plants, and ∼3.8, ∼4.0 and ∼4.3-fold up-regulation was observed in roots of 500 mM NaCl treated plants at 1, 4 and 8DAS, respectively. Further, Ca^2+^ levels were slightly increased in roots of 300 mM NaCl treated plants at 1, 4 and 8DAS. While, in roots of 500 mM NaCl treated plants, Ca^2+^ levels were increased significantly at 1 and 4DAS, while these levels were unchanged at 8DAS.

### Visualization of Na^+^ ion specific fluorescence in leaves and roots of P. pinnata

To investigate the pattern of Na^+^ ion accumulation in leaves and roots of *P. pinnata*, a cell permeable Na^+^ specific fluorescent probe CoroNa-Green AM was used (Supplementary Figure 3A-J and Supplementary Figure 4A-T). Leaf sections of control plants were showed no strong Na^+^ specific fluorescence signal at 1, 4 and 8DAS. Similarly, leaf sections of 300 mM NaCl treated plants were also showed no strong Na^+^ specific fluorescence signal at 1 and 4DAS, whereas a Na^+^ specific green florescence was observed in apoplasmic region at 8DAS. Conversely, we could observe a consistent increase in the Na^+^ fluorescence signal in apoplasmic regions in leaf sections of 500 mM NaCl treated plants at 1, 4 and 8DAS. Further, roots were followed a different pattern of fluorescence in both control and salt treated plants. We could observe no strong fluorescence signal in the cortical apoplasmic regions of control root sections at 1, 4 and 8DAS. However, we could observe a strong fluorescence signal in the cortical apoplasmic regions dose dependently in both 300 and 500 mM NaCl treated root sections at 1, 4 and 8DAS.

### Salinity-induced phytohormonal changes in leaves and roots of P. pinnata under progressive salt stress

Upon salt stress, Pongamia accumulated various hormones tissue specifically and time dependently. Accordingly, we quantified phytohormones by using LC-MS analysis. All seven hormones zeatin, IAA, IBA, MeJA, SA, ABA, and JA were showed a significant increase in leaves of 300 and 500 mM NaCl treated plants at 1, 4 and 8DAS (Figure 1B-D). As shown in the Figure 1B-D, the hormone accumulation pattern was changed with the treatment time in leaves of 300 mM NaCl plants. At 1DAS, six hormones zeatin, IAA, IBA, MeJA, ABA, and JA were showed significant increase in leaves of 300 mM NaCl treated plants, while SA levels did not change significantly. Hormones such as zeatin, IAA, MeJA and JA showed significant up-regulation at 4DAS. IAA, ABA and JA levels were significantly enhanced in leaves of 300 mM NaCl treated plants at 8DAS. Moreover, in roots, the phytohormone levels varied with salt concentration and treatment time. Zeatin, IBA and MeJA levels increased significantly in roots of 300 mM NaCl treated plants at 1DAS. Similarly, zeatin and ABA levels showed significant up-regulation in 500 mM NaCl treated plants at 1DAS (Figure 1E). At 4DAS, only zeatin showed significant increase by ∼3.7-fold in roots of 300 mM NaCl treated plants. In response to 500 mM NaCl treatment, MeJA and JA levels showed ∼3.3 and 2.0-fold up-regulation in roots at 4DAS respectively (Figure 1F). At 8DAS, zeatin and IAA levels were increased by ∼2.2 and 2.3-fold in roots of 300 mM NaCl treated plants respectively (Figure 1G). In response to 500 mM NaCl stress, only IBA levels showed significant increase by ∼3.0-fold at 8DAS.

Correlation studies among hormones are essential to explore hormonal interactions under salt stress. Accordingly, correlation studies were performed based on stress treatment time. Each stress time point was grouped individually in the form of a correlation cluster matrix to show the interactive responses of each individual hormone under salt stress. We observed significant correlation between JAs and IAA over time in leaves of 300 mM NaCl treated plants (Figure 2A-C). Similarly, JA showed significant correlation with MeJA in leaves of 500 mM NaCl treated plants across all-time points (Figure 2D-F). Jasmonates (JA and MeJA) and SA showed significant interaction with other hormones in roots of 300 and 500 mM NaCl at all-time points (Figure 2G-L).

**Fig. 2.**
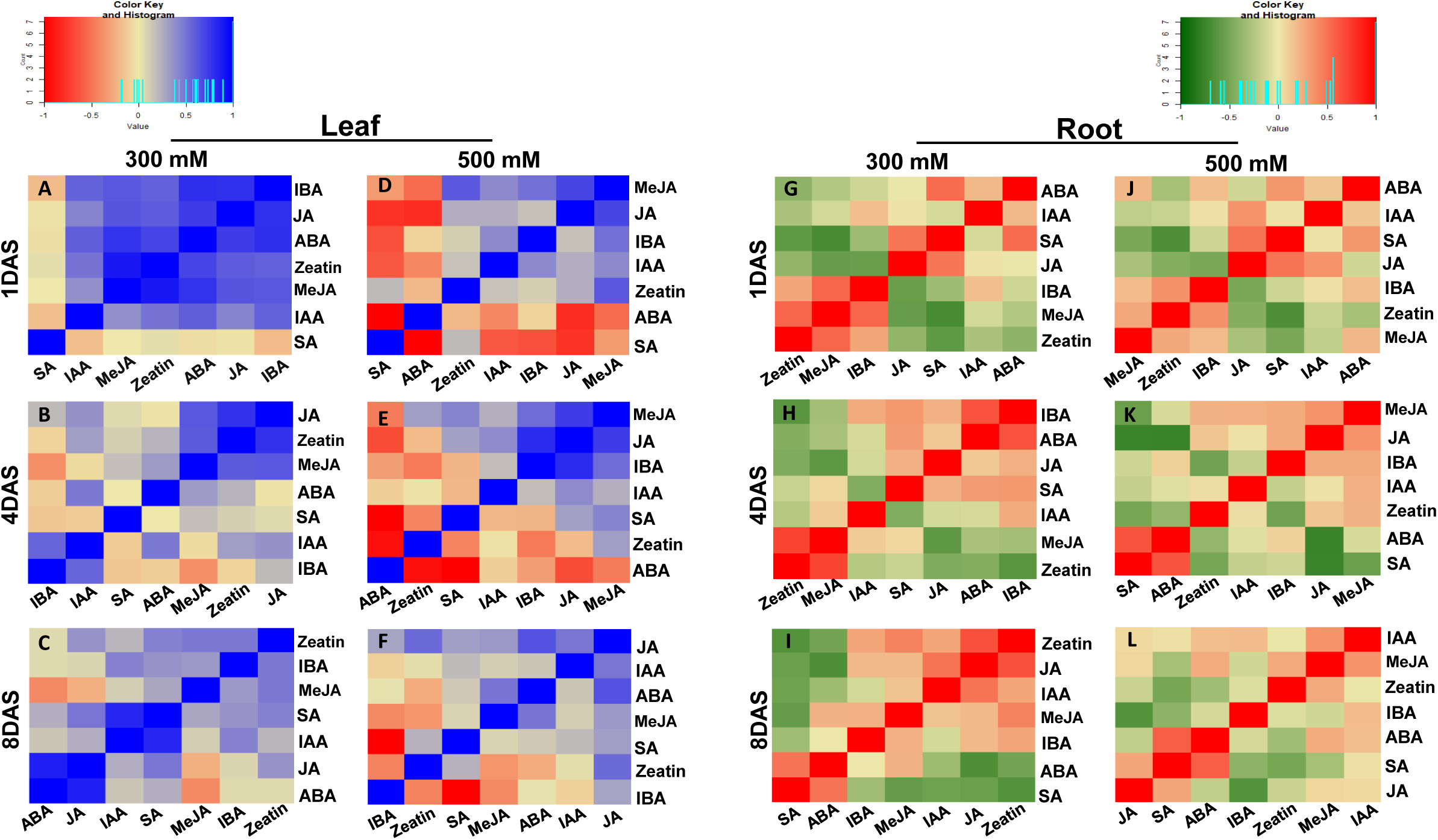
Hierarchical cluster analysis and heat map of hormone-hormone correlation matrix in *P. pinnata* under different NaCl treatments. Each correlation value (based on Pearson correlation coefficient) corresponds to average of six biological replicates. HAC analysis performed among the hormones at each individual time points (A and D) 1DAS, (B and E) 4DAS, (C and F) 8DAS in leaves as well as (G and J) 1DAS, (H and K) 4DAS, (I and L) 8DAS in roots of 300 and 500 mM NaCl treated plants. The colour key and histogram show degree of correlation.

### Temporal metabolite profiling in leaves and roots of Pongamia pinnata under salt stress

The relative levels of metabolites in Pongamia under salt stress conditions were quantified by using gas chromatography-mass spectrometry (GC-MS) (Supplementary Figures 5 and 6). A total of 71 metabolites were identified in both leaves (Supplementary Table 1) and roots (Supplementary Table 2), which are related to 12 carbohydrate metabolism, 22 organic acid metabolism, 15 amino acid metabolism, 6 cyclitol/ polyol metabolism,6 fatty acid metabolism and 10 other miscellaneous metabolites.

In response to salinity stress, leaves accumulated a set of primary metabolites related to various metabolic pathways such as amino acid metabolism, carbohydrate metabolism, organic acid metabolism, polyol and fatty acid metabolism (Supplementary Figure 5). Upon salt stress exposure, leaves accumulated various metabolites, which includes Suc, Fru, Man, pinitol, mannitol, myo-inositol, Phe, Val, butanoate, coumarate and myristate showed significant increase in both 300 and 500 mM NaCl treated plants at 1, 4 and 8DAS. Importantly, mannitol levels showed significant up-regulation by ∼12 fold in leaves of 300 and 500 mM NaCl treated plants at 1, 4 and 8DAS. With increasing treatment time, Pongamia accumulated several metabolites related to cell wall synthesis and TCA cycle, which include Gal, Xyl, pGlc, pFru, Leu, fumarate, succinate and glutarate in leaves of 300 and 500 mM NaCl treated plants at 8DAS. Additionally, several other metabolites were increased marginally in related to glycerol pathway, organic acid synthesis, amino acid metabolism and GABA shunt in leaves of treated plants across all-time points.

In response to salt stress, roots accumulated a set of primary metabolites related to various metabolic pathways such as amino acid metabolism, carbohydrate metabolism, amino acid metabolism, polyol and fatty acid metabolism (Supplementary Figure 6). Additionally, roots of salt treated plants also accumulated various metabolites related to cell-wall carbohydrates, carbohydrate alcohol, polyol, amino acid and fatty acid metabolism, which includes Gal, mannitol, NAG, myristate, laurate, palimitate, phytol and methyl-succinate showed significant increase in 300 and 500 mM NaCl treated plants at 1, 4 and 8DAS. Importantly, similar to leaves, mannitol levels showed significant up-regulation by ∼12 fold in roots of 300 and 500 mM NaCl treated plants at 1, 4 and 8DAS. In addition, Pongamia leaves accumulated several metabolites related to cell wall synthesis, polyols and organic acids, which includeMan, Fru, pGlc, pTag, pinitol, Val, Leu, glycerol, threonate and benzoate in roots of 300 and 500 mM NaCl treated plants at 8DAS. Several other metabolites also increased marginally relating to fatty acid, organic acid synthesis, amino acid metabolism and GABA shunt in leaves of treated plants across all-time points.

### Correlation based clustering among metabolites in leave and roots P. pinnata under salt stress

In order to explore the relationship among the metabolites in Pongamia subjected to salt stress, HAC (hierarchical cluster) analysis was performed on a total 60,492 elements of two different tissues leaf (30,246) and root (30,246) which include two different treatments 300 mM NaCl (15,123), 500 mM NaCl (15,123) and three time points 1, 4 and 8DAS (71 × 71 metabolite; 5041) (Figures 3 and 4). Correlation analysis was performed with Pearson correlation coefficients for each metabolite pair (Moing et al., 2011; Sánchez et al., 2012; Lombardo et al., 2011). The correlation values were clustered into six groups based on treatment time e.g. leaves of 300 mM NaCl treatment at 1DAS clustered into a group (Figure 3A and D; order of metabolite names and abbreviations were given in the Supplementary Table 3), leaves of 300 mM NaCl treatment at 4DAS clustered into a group (Figure 3B and E; Supplementary Table 3) and leaves of 300 mM NaCl treatment at 8DAS clustered into another group (Figure 3C and F; Supplementary Table 3). In order to simplify the graphics, only strong correlations (r≥0.6) were showed separately in the tables. Similarly, leaves of 500 mM NaCl at 1DAS clustered into one group (Figure 3G and J; order of metabolite names and abbreviations were given in the Supplementary Table 4), leaves of 500 mM NaCl at 4DAS clustered into one group (Figure 3H and K; Supplementary Table 4) and leaves of 500 mM NaCl at 8DAS clustered into one group (Figure 3I and L; Supplementary Table 4). In leaves of 300 mM NaCl treated plants, we observed high correlations within the organic acid group (e.g. CA:CAA:NA, 5HPip:Pip) and polyols (MI:MT). Although a significant correlation was not observed within the amino acid group, these levels showed strong interaction with organic acids (Asp:CA:NA:CAA, Asn:Pip:5HPip, GABA:MA) in leaves of 300 mM NaCl treated plants. Additionally, a strong positive correlation was observed between several metabolites such as fructose, myristate, mannitol and myo-inositol in leaves of 300 mM NaCl treated plants. In 500 mM NaCl treated plants, a significant correlation was observed within the amino acids group (Asp:Glu:Oxopro, Asn:Hse, Leu:Phe), organic acid group (e.g. 5HPip:CA:CAA, IBua:oC) and carbohydrate group (Xyl:pGlc, Man:pFru) at all-time points. Carbohydrates also showed significant interaction with other metabolites (Suc:Val, Xyl:SA:Pi, pGlc:SA) in 500 mM NaCl treated plants. Additionally, several metabolites showed a significant interaction within each other (e.g. Asp:Oxopro:Glu:5Hpip:CA:CAA:Grl-g, Man:pFru:MI:MT:oC:IBu, Asn:Hse:Pip) across all time points.

**Fig. 3.**
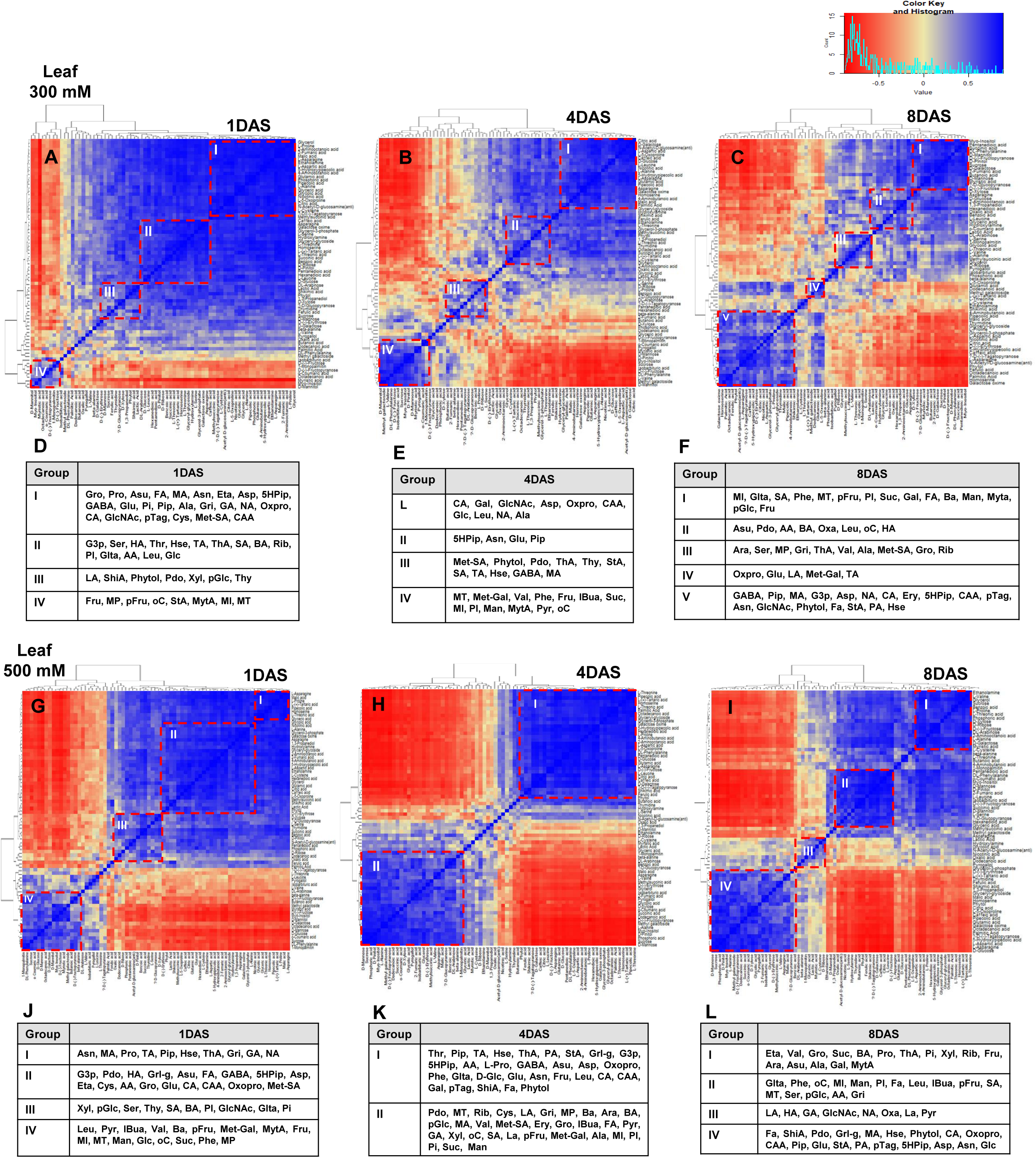
Heat map of hierarchical clustering of metabolite-metabolite correlations in leaves and roots of 300 mM NaCl treated *P. pinnata* under salinity stress. Each correlation value (based on Pearson correlation coefficient) corresponds to average of six biological replicates. HAC analysis performed among the metabolite at each individual time points (A) 1DAS, (B) 4DAS and (C) 8DAS in leaves of 300 mM NaCl treated plants. (D, E and F) Detailed view of positively correlated metabolite correlations was shown in the form of table. HAC analysis performed among the metabolite at each individual time points (G) 1DAS, (H) 4DAS and (I) 8DAS in leaves of 500 mM NaCl treated plants. (J, K and L) Detailed view of positively correlated metabolite correlations was shown in the form of table. The colour key and histogram show degree of correlation.

**Fig. 4.**
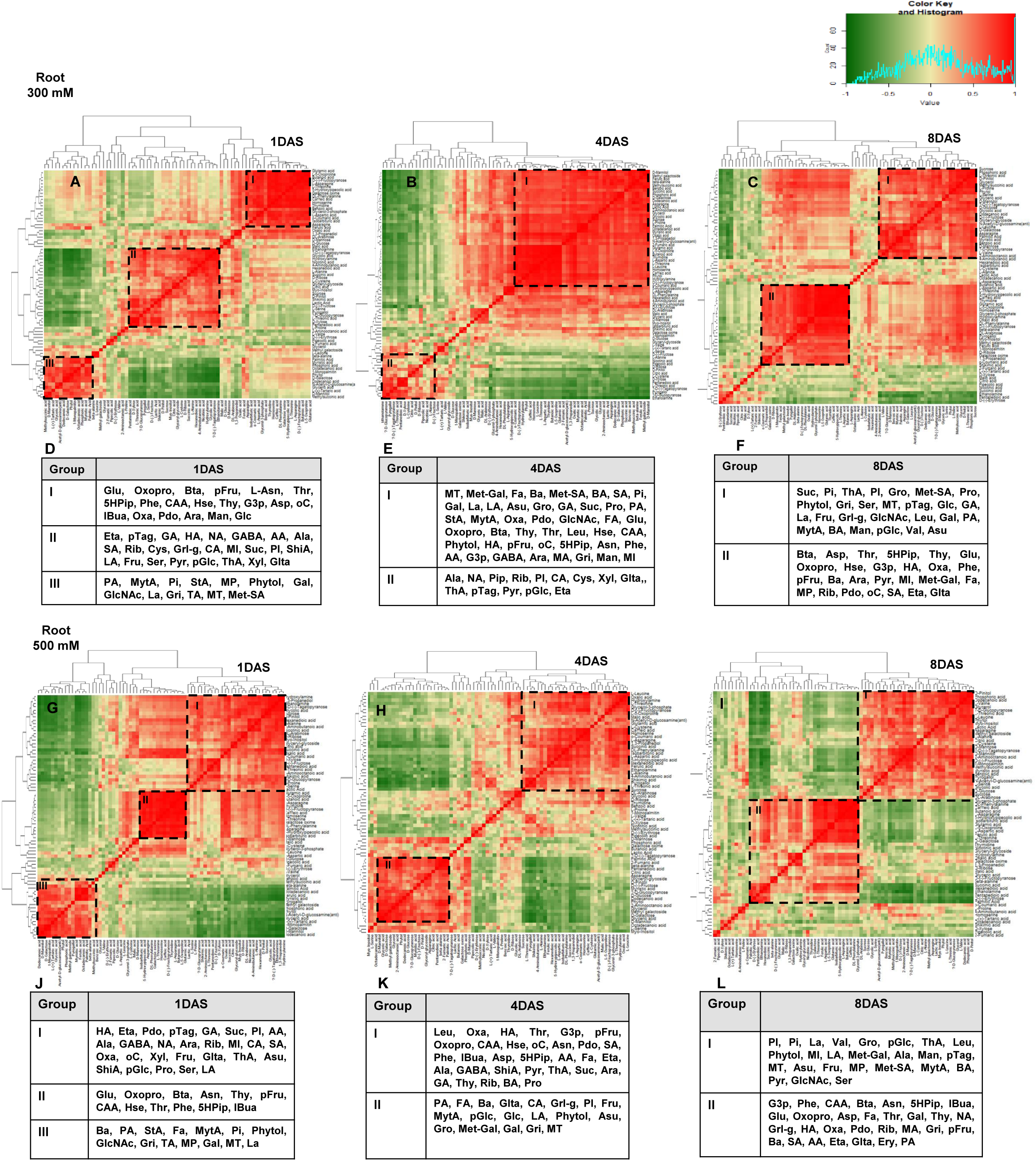
Heat map of hierarchical clustering of metabolite-metabolite correlations in roots of 300 mM NaCl treated *P. pinnata* under salinity stress. Each correlation value (based on Pearson correlation coefficient) corresponds to average of six biological replicates. HAC analysis was performed among the metabolite at each individual time points (A) 1DAS, (B) 4DAS and (C) 8DAS in roots of 300 mM NaCl treated plants. (D, E and F) Detailed view of positively correlated metabolite correlations was shown in the form of table. HAC analysis was performed among the metabolite at each individual time points (G) 1DAS, (H) 4DAS and (I) 8DAS in roots of 500 mM NaCl treated plants. (J, K and L) Detailed view of positively correlated metabolite correlations was shown in the form of table. The colour key and histogram show degree of correlation.

In the root dataset, the correlations among the metabolites were less pronounced when compared with leaf datasets. In roots of 300 mM NaCl treated plants, most of the metabolite levels showed significant correlation with each other (e.g. Glu:Oxopro:Thr:Phe:Hse:Ara:5Hpip:oC:Oxa:Pdo, PA:MytA:Gal:Gri:Met-SA:MT:Pi:Phytol:GlcNAc,pGlc:ThA:PI:pTag, Rib:Pyr:Glta:Eta, Suc:GA:Fru, HA:MI:SA) (Figure 4A-F; names and abbreviations were given in the Supplementary Table 5). In 500 mM NaCl treated plants, a strong positive correlation was observed within the organic acid group (SA:Oxa) and carbohydrate group (pGlc:Fru) across all the time points (Figure 4G-L; names and abbreviations were given in the Supplementary Table 6). In addition, several metabolites were associated significantly within each other (e.g. Oxopro:Asn:Thr:Phe:CAA:5Hpip:IBua:pFru:Thy, Ala:ThA, LA:Asu:PI, Rib:AA, HA:Pdo:Eta, Ba:Gal:Gri:PA, MytA:MT:Phytol) in 500 mM NaCl treated plants at all-time points.

### Correlation based clustering between hormones and metabolites in leaves and roots of P. pinnata under salt stress

Figure 5 shows the interaction between hormones and metabolites. In leaves, hormones including ABA, IBA, MeJA, zeatin and JA showed a positive correlation with MytA in leaves of 300 mM NaCl treated plants at 1DAS (Figure 5A). Hormones JA, MeJA and zeatin showed significant interaction with metabolites such as amino acids, carbohydrates, and organic acids in leaves of 300 mM NaCl treated plants at 4DAS (Figure 5B). In addition, JA and MeJA also showed strong association with carbohydrates and amino acids in leaves of 300 mM NaCl treated plants at 4DAS. At 8DAS, JA and ABA showed significant correlation with several metabolites MytA, Suc, Gal, Man and FA, while ABA and JA were showed significant interaction with metabolites pTag, organic acid group (NA, Pip, MA, 5Hpip, CAA), amino acids group (Hse, Asp, Asn), StA, and other metabolites (G3p, NAG, Grl-g) (Figure 5C). In leaves of 500 mM NaCl treated plants, we could observe more interaction between hormones and metabolites including MeJA:IBA:zeatin:JA:IAA::carbohydrates, JA:MeJA:IAA:zeatin::organic acids, ABA:IBA:IAA:zeatin::amino acids, JA:IBA:IAA:SA:zeatin::polyols and fatty acids at 1DAS (Figure 5D). Increased interactions were also observed between hormones and metabolites namely JA:MeJA:zeatin::carbohydrates and cell-wallcarbohydrates, JA:MeJA:IBA:zeatin::organic acids, MeJA:IBA:IAA::amino acids, JA:MeJA:zeatin::polyols at 4DAS (Figure 5E). We recorded a significant positive correlation between hormones and metabolites namely MeJA:JA:ABA:SA::carbohydrates and cell-wall carbohydrates, JA:ABA:MeJA:SA::organic acids, JA:ABA:MeJA:IBA:IAA::amino acids, JA:ABA::polyols, fatty acids and JA:ABA:MeJA:IBA:zeatin::fatty acids in leaves of 500 mM NaCl treated plants at 8DAS (Figure 5F).

**Fig. 5.**
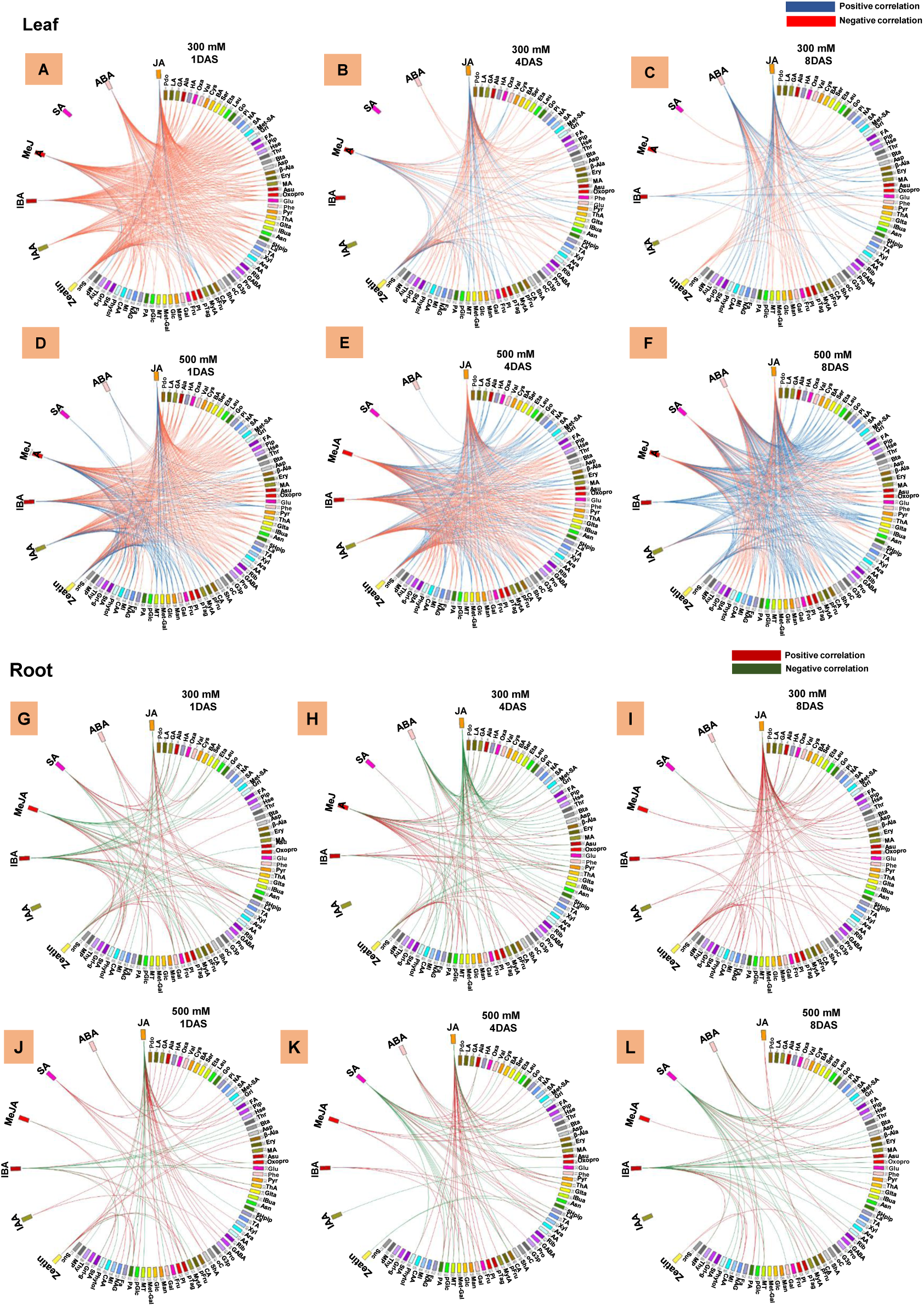
Circos plots showing correlations in the hormone-metabolite interactions in *P. pinnata* under different NaCl treatments. All 71 metabolites and 7 hormones were identified in the circle (order of metabolites mentioned in the Supplementary Table 3, 4, 5 and 6). Each correlation value (based on Pearson correlation coefficient) corresponds to average of six biological replicates and analysis performed between the hormones-metabolites at each individual time points (A and D) 1DAS, (B and E) 4DAS, (C and F) 8DAS in leaves as well as (H and K) 1DAS, (I and L) 4DAS, (J and M) 8DAS in roots of 300 and 500 mM NaCl treated plants. Ribbon colour corresponds to degree of correlation (in leaves: blue (+ve correlation) and red (−ve correlation) and in roots: red (+ve correlation) and green (−ve correlation).

In roots, significant associations between hormones and metabolites were including JA:SA:ABA:MeJA:IBA:zeatin::carbohydrates, JA:MeJA:SA:ABA:IAA:SA::organic acids and MeJA:IBA::fatty acids in 300 mM NaCl treated plants at 1DAS (Figure 5G). At 4DAS, more positive interactions were seen between hormones and metabolites namely JA:ABA:ABA:SA:IBA::cell-wall carbohydrates, MeJA:JA:zeatin::polyols and MeJA:zeatin::fatty acids (Figure 5H). At 8DAS, a strong interaction was noticed between hormones and metabolites (JA:MeJA:ABA:zeatin::organic acids, JA:IBA:zeatin::amino acidand JA:zeatin::polyols) (Figure 5I). As the salinity progressed from day 1 to day 8, the positive interactions decreased in roots of 500 mM NaCl treated plants (Figure 5J-L). In roots of 500 mM NaCl treated plants, significant interaction was between hormones and metabolites namely ABA:SA:IBA:zeatin::organic acids at 1DAS, JA:SA:IBA::cell-wall carbohydrates, ABA:SA:JA:MeJA::organic acids at 4DAS, and ABA:SA::cell-wall carbohydrates, ABA:MeJA::organic acid at 8DAS respectively.

### Differential expression of genes associated with ion transport and membrane potential in leaves and roots of P. pinnata

To assess the effect of salinity stress on the gene expression profile of Pongamia, we monitored the expression profile of several key salt-responsive genes including SOS pathway components (SOS1, SOS2 and SOS3), transporters (NHX1, HKT1:1, CLC1, V-CHX1, CCX1, V-H^+^ATPaseB subunit, V-H^+^ATPaseE subunit, PM-H^+^ATPase1, PM-H^+^ATPase4.1, PM-H^+^ATPase4.1-like, CNGC5 and CNGC17), and calcium-dependent protein kinases (CDPK3 and CDPK32) (Figure 6).

**Fig. 6.**
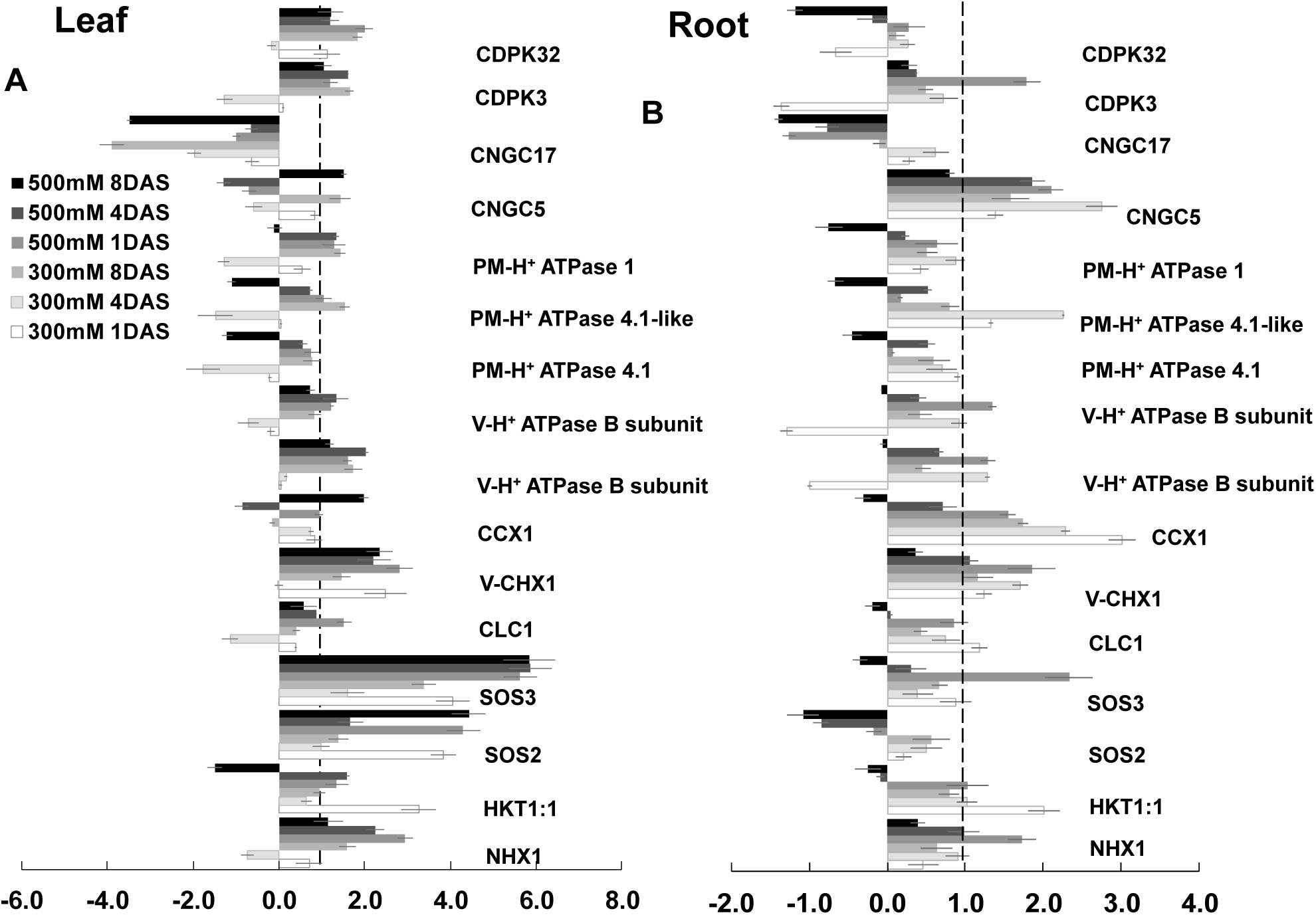
Relative mRNA expression levels of transporters in leaves of *P. pinnata* under salt stress conditions. Log_2_ fold changes of NHX1, HKT1:1, SOS2, SOS3, CLC1, V-CHX1, CCX1, V-H^+^ATPaseB subunit, V-H^+^ATPaseE subunit, PM-H^+^ATP4.1, PM-H^+^ATP4.1-like, PM-H^+^ATPase1, CNGC5, CNGC17, CDPK3, and, CDPK32 in leaves (A) and roots (B) of salt-treated plants of *P. pinnata* at 1, 4 and 8 DAS respectively when compared to their corresponding controls. Error bar represents the mean ± SD (n=6).

Sodium proton exchanger 1 (NHX1) was significantly up-regulated by ∼2.6 and ∼3.6-fold in leaves of 500 mM NaCl treated plants at 1 and 4DAS respectively, while these levels remained unchanged in leaves of 300 mM NaCl treated plants at all-time points and 500 mM NaCl at 8DAS respectively (Figure 6A). Further, the NHX1 gene expression was significantly increased only in 500 mM NaCl treated plants at 1DAS (Figure 6B). Interestingly, high affinity transporter 1:1 (HKT1:1) levels significantly increased by ∼3.6 and ∼2.0-folds in both leaves and roots of 300 mM NaCl treated plants at 1DAS. The SOS (Salt Overly Sensitive) 2 levels showed increase/ decrease in leaves of 300 and 500 mM NaCl treated plants, while these levels were unchanged in the roots of salt treated plants. SOS3 levels showed a significant up-regulation by ∼3.9, ∼2.0, and ∼3.7-fold in leaves of 300 mM NaCl treated plants as well as ∼5.9, ∼6.0 and ∼6.0-fold increase was observed in laves of 500 mM NaCl treated plants respectively. However, we did not observe much changes in expression levels of chloride channel 1 (CLC1) in both leaves and roots of salt treated plants. The vacuolar cation proton exchanger 1 (V-CHX1) levels were marginally induced in both leaves and roots of salt treated plants. Under salt stress, cation calcium exchanger 1 (CCX1) levels were induced significantly by ∼2.0-foldin 300 mM NaCl treated plants only at 8DAS, while these levels significantly increased by ∼3.0, ∼2.5 and 1.9-fold in roots of 300 mM NaCl treated plants, though these levels remained unchanged in 500 mM NaCl treated plants. Vacuolar proton ATPaseB subunit (V-H^+^ATPaseB subunit) and V-H^+^ATPaseE subunit expression levels were slightly induced in both leaves and roots of salt treated plants. We also monitored expression levels of three plasma membrane proton ATPase (PM-H^+^ATPase) isoforms PM-H^+^ATPase1, 4.1, 4.1-like in both leaves and roots of salt treated plants. The expression levels of PM-H^+^ATPase1, 4.1, and 4.1-like genes were similar to those of control levels in leaves of salt treated plants, while PM-H^+^ATPase4.1-like levels were marginally increased in roots of 300 mM NaCl treated plants only at 4DAS. Cyclic nucleotide-gated ion channel 5 (CNGC5) levels were decreased or increased in leaves of treated plants while these levels were significantly induced in roots of treated plants at 1, 4 and 8DAS. Furthermore, the expression level of CNGC17 gene was decreased/ unchanged in both leaves and roots of treated plants. Calcium-dependent protein kinase3 (CDPK3) and CDPK32 levels were increased/ unchanged in both leaves and roots salt treated plants.

Gene primers for NHX3, NHX6, NHX6-like, SOS1, SOS1-like, H^+^-ATPase4, H^+^-ATPase4-like, CNGC17-like, and V-H^+^ppase exhibited with multiple banding patterns their expression pattern significantly varied in leaves and roots of both control and salt-treated plants (given in Supplementary Figures 7 and 8, primers were given in the Supplementary Table 7).

### Differential expression of genes associated with ROS homeostasis and cell wall modifications in leaves and roots P. pinnata

We monitored the expression levels of several key salt-responsive genes including antioxidant enzymes (CAT4, FeSOD, MnSOD, Cu/ZnSOD1, and Cu/ZnSOD1-like) and cell wall thickening enzymes (POD, PAL1, PAL2, CAD2, and CAD6) (Supplementary Figures 9A, 10 and 11).

Among the antioxidant genes CAT4 expression remained unchanged in leaves throughout the salt stress treatment and showed down-regulation at 8 DAS in both 300 and 500 mM salt stress. In leaves, different isoforms of SODs showed gradual and significant up-regulation under 300 mM salt treatment from 1 to 8DAS. However, under 500 mM NaCl treatment, the SOD isoforms showed significant up-regulation only till 4DAS, but declined at prolonged stress treatment at 8DAS. Among the cell-wall strengthening genes, only POD showed a strong up-regulation in leaves of 300 and 500 mM NaCl treated plants at all-points, while the other genes (PAL1, PAL2, CAD1 and CAD2) did not show any significant induction. CAD 6 showed significant increase in 300 mM NaCl treatment at 8DAS, while these levels were significantly down-regulated in 300 mM NaCl 1 and 4DAS as well as in 500 mM NaCl treated plants at 1, 4, and 8DAS.

In roots, the expression levels CAT4 were substantially up-regulated in 300 and 500 mM NaCl treated plants at 1, 4 and 8DAS respectively (Supplementary Figures 9B, 10 and 11). Among the antioxidant genes, CAT4 was significantly up-regulated only at 1DAS of 500 mM salt treatment. FeSOD also increased at all-time points in 300 mM salt treated plants and in 500 mM NaCl treatment at 4 and 8DAS. However, the expression levels of all other genes including MnSOD, Cu/ZnSOD1, Cu/ZnSOD1-like SOD, PAL1, 2, and CAD1, 2 and 6 were slightly changed in roots of salt treated plants.

## Discussion

### Sustained photosynthesis parameters, Ca^2+^ signalling pathways and apoplasmic Na^+^ sequestration in the roots are the possible salt adaptive mechanisms in Pongamia

Pongamia exhibited significant tolerance to high salinity (500 mM NaCl (∼3% NaCl) like true halophytes and mangroves by exhibiting sustained leaf morphology and insignificant alterations in the photosynthetic machinery (Marriboina et al., 2017). The reduced LRWC and unchanged RRWC indicate that the roots were able to process salt solution (Marriboina Attipalli, 2020a). The results regarding Na^+^ and Cl^-^ ions accumulation are rather interesting. High levels of Na^+^ and Cl^-^ ions were detected in roots of treated plants suggesting that the roots may act as potential sink for excessive Na^+^ and Cl^-^ ions deposition, inhibiting their translocation to shoot and alleviate the negative effects on actively dividing and photosynthesizing cells (Baetz et al., 2016; Peng et al., 2016; Rahneshan et al., 2018; Wu et al., 2018). Intriguingly, Ca^2+^ levels were also increased with increasing Na^+^ levels in both leaves and roots of salt treated plants, which suggest that might possible trigger of vaculoar Ca^2+^ reserves to ameliorate the Na^+^ toxicity (Saleh Plieth, 2013). Ca^2+^ was found to be involved in propagation of long distance signalling (root-to-shoot) under salt stress conditions (Choi et al., 2014).

Our previous fluorescence studies demonstrated that Pongamia roots act as an effective sequester of Na^+^ ions (Marriboina Attipalli, 2020a). In the present study, the apoplastic/ cell wall Na^+^ specific fluorescence intensity was increased with treatment time in both 300 and 500 mM NaCl treated plants indicating that there was an apoplastic Na^+^ sequestration (Gonzalez et al., 2012; Anower et al., 2017). The carboxylic residues of pectin in the cell wall may primarily serve as cation-binding matrix for Na^+^ ion, contributing to apoplastic Na^+^ sequestration (Gonzalez et al., 2012; Marriboina et al., 2017). The patterns of high apoplastic and vaculoar Na^+^ contents might lead to lower Na^+^ ion content in the cytosol (Marriboina Attipalli, 2020a), which may facilitate the protection of cytosolic enzymes from sodium toxicity (Wu, 2018).

### The consequence of salt-induced phytohormonal disturbance in leaves and roots of Pongamia

Developmental plasticity under stress conditions largely depends upon the interactions between hormones, which regulate stress-adaptation responses and developmental processes. In this study, both leaves and roots of salt treated plants showed a significant diversity in hormone profile and their correlation patterns at all-time points. A significant correlation was observed among all hormones due to initial exposure of 300 mM NaCl stress in leaves. The rise in all hormones and correlation may be beneficial for the plant to maintain the growth under salt stress conditions (Sahoo et al., 2014; Fahad et al., 2015). Moreover, increased levels of zeatin in both leaves ad roots of salt treated plants improved RRWC and stress-induced growth under salt stress conditions (Ghanem et al., 2011; Nishiyama et al., 2011; Wu et al., 2014; Melo et al., 2016). A strong correlation was also observed between zeatin and JAs in leaves and roots of 300 mM NaCl treated plants. The interactive influence of cytokinin may substantiate JAs negative impact to promote plant survival under extreme saline conditions. The synergistic interaction between zeatin and JAs may also enhance the salt induced-vasculature in roots to enhance water uptake, which is also supported by the well maintained RRWC in roots of treated plants (Supplementary Figure 2D) (Ueda Kato, 1982; Nitschke et al., 2016; Jang et al., 2017).

IAA levels were significantly increased in leaves of salt treated plants at all-time points, indicating that IAA might play an important role in Pongamia salt tolerance. Transgenic poplar plants, overexpressing *AtYUCCA6*gene associated with increased levels of auxin, showed delayed chlorophyll degradation and leaf senescence (Kim et al., 2012). The increased levels of IAA and IBA may due to tissue damage or cell lysis. However, our fluorescence studies clearly suggest the viable status of cells and tissue integrity. Therefore, increased IAA levels in Pongamia might help in maintaining “stay-green” trait and steady levels of chlorophyll pigments under salt stress (Kim et al., 2012). Likewise, IBA levels also showed significant increase in both leaves and roots of salt treated plants, suggesting that enhanced IBA levels may also play a role in acquiring the stress-induced protective architectural changes in Pongamia (Tognetti et al., 2010). In addition, correlation studies revealed that IBA showed good interaction with IAA in leaves of 300 mM NaCl treated plants. It was evident that the plants deficient in both IAA and IBA genes showed defective plant growth and development (Spiess et al., 2014). Auxins also showed a strong association with JAs (JA and MeJA) in leaves and roots of salt treated plants, which might involve in promoting salt-induced growth and tissue integrity during salt stress (Cai et al., 2014; Fattorini et al., 2018; Ishimaru et al., 2018).

A significant increase in JAs was detected in both leaves of salt treated plants. However, in roots, JA levels were maintained low till 4DAS, and restored to control levels at 8DAS. Conversely, MeJA levels were maintained high till 4DAS, while these levels returned to control levels at 8DAS. The results indicates that the two different forms of JAs are presumably interchangeable and might share common signal transduction pathway in Pongamia during salt stress (Diallo et al., 2014; Mitra Baldwin, 2014; Cao et al., 2016; Li et al., 2017). An exogenous application of JAs reduced shoot growth, enhanced water uptake and cell wall synthesis in certain crop species (Kang et al., 2005; Uddin et al., 2013; Shahzad et al., 2015; Tavallali Karimi, 2019). Previous studies suggest that the increased JAs level alleviate the toxic effects of salt stress by lowering the Na^+^ and Cl^-^ ions accumulation across the plant (Shahzad et al., 2015). Correlation studies revealed that JAs showed a strong correlation with ABA in leaves of treated plants. The interactions between JAs and ABA may induce stomatal closure by triggering the stress-induced signalling pathways in guard cells, preventing water loss from the leaves (Munemasa et al., 2011; Wang et al., 2016; Yang et al., 2018) (Figure 7).

**Fig. 7.**
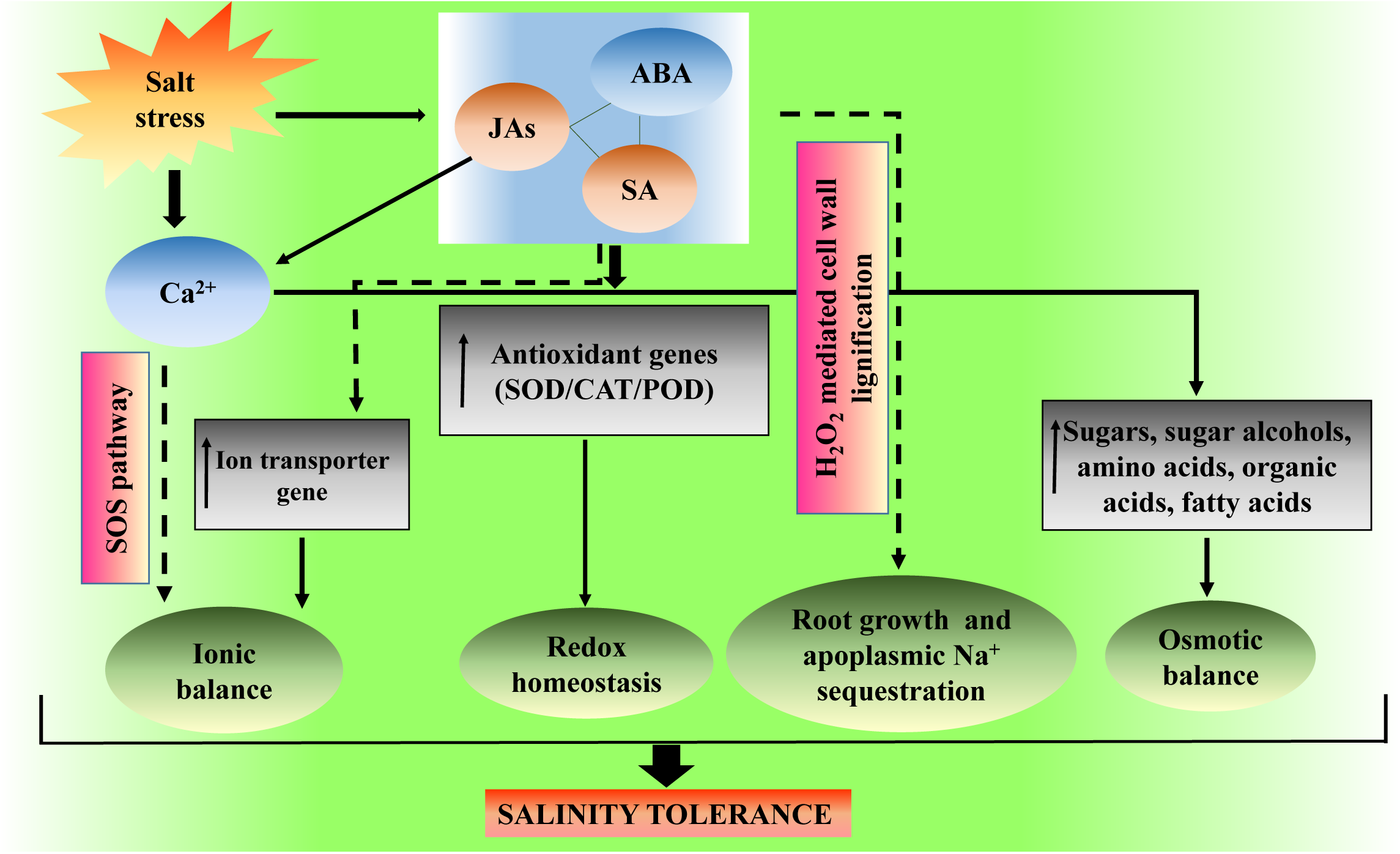
Our proposed model based to show the mechanism of high salinity tolerance in *Pongamia pinnata*.

ABA levels were significantly increased in leaves of salt treated plants which might limit the stomatal conductance, water content, transpiration rate and CER by closing the stomatal apparatus (Skorupa et al., 2019). The observed reduction in ABA accumulation in roots might be due to ABA transfer from root to shoot or ABA exudation from the roots (Shi et al., 2015). The prolonged maintenance of higher ABA levels negatively impacts the plant growth, while transient increase helps in mitigation of salt stress by enhancing the stress responsive genes (Shi et al., 2015). Further, reduced ABA content may favour in maintaining RWC by regulating aquaporin proteins (Shi et al., 2015). The correlation studies revealed that ABA showed a strong interaction with SA in roots of salt treated plants, which improves plant growth under saline conditions, albeit the signalling mechanism is still unclear (Devinar et al., 2013) (Figure 7).

SA levels in roots were maintained little low at 1DAS and maintained high at 4DAS, while these levels returned to control levels at 8DAS. Exogenous application of SA on plants showed an improvement in LRWC and ROS homeostasis under salt stress (Jayakannan et al., 2013; Husen et al., 2018). The rise in SA levels might protect the leaves from salt injury by inhibiting necrosis signalling pathways and also regulate the leaf turgor by accumulating carbohydrate polyols (mannitol, pinitol, and myo-inositol) (Husen et al., 2018). The observed reduction in SA levels in roots may be due to the transportation of SA from root to shoot under salt stress conditions (Xu et al., 2017). Correlation studies revealed a positive correlation between SA and IAA in leaves of salt treated plants might help in maintaining the leaf cell extensibility to promote better plant growth under salt stress (Formentin et al., 2018; Shaki et al., 2019).

### Enhanced cell-wall carbohydrates, carbohydrate alcohols, organic and fatty acids maintain cellular osmotic balance under salt stress

Pongamia exhibited significant changes in the metabolite level including carbohydrates, amino acids, organic acids, polyol, and fatty acids. Interestingly, the rise in mannitol level (∼12 fold), observed in both leaves and roots of salt treated plants at 8DAS, suggest that mannitol may serve as a major osmoticum in Pongamia. The increased levels of carbohydrate alcohols not only regulates the cellular osmotic potential but also provides non-enzymatic ROS scavenging protection under abiotic stress conditions (Abebe et al., 2003; Hossain et al., 2017; Dumschott et al., 2019). High levels of carbohydrates were recorded in both leaves and roots of treated plants at 8DAS. These carbohydrates are known to contribute to secondary call wall synthesis, while their accumulation alters the cell wall composition in Pongamia (Gilbert et al., 2009; Geilfus et al., 2017; Zhao et al., 2019). The rise in carbohydrates level may serve as osmolytes and immediate source of energy for a cell (Abdallaha et al., 2016).

The metabolic profile also indicate that enhanced levels of free amino acids such as β-Ala, Val, Leu, Ala, Thr, Cys and Phe in leaves and roots of treated plants at 8DAS. The increased levels of these free amino acids may provide continue nutrient and water uptake to support plant growth under salt stress condition and also contribute to tolerance by regulating several biological processes including biosynthesis of cell wall components, to protect the membrane protein integrity as well as stability of cellular macro-structures ((Cao et al., 2017; Nasir et al., 2010). Interestingly, the accumulation of serine and glycolate are active components of photorespiration, which play a crucial role in protecting the photosynthestic apparatus by limiting the deposition of toxic photo-inhibitory metabolites (Hossainet al., 2017). Lactic acid accumulation may involve in the regulation of cytoplasmic pH, and production of pyruvate to maintain glycolysis under salt stress condition (Felle et al., 2005; Hossain et al., 2017).

Increased accumulation of saturated fatty acids might protect the membrane fluidity to protect the cell and cellular organelle from Na^+^ toxicity and ion leakage (Shu et al., 2015; Atikij et al., 2019). Our results also showed an increase in coumarate and benzoate in both leaves and roots, while significant increase was observed in caffieate and ferulate levels only in roots of salt treated plants. Increased levels of these metabolites could be beneficial to the plants for lignin biosynthesis as well as salicylic acid production, which may play defensive role under salinity stress (Chen et al., 2019). An increase in the accumulation of free amino acids Thr, Ala, Cys, and Leu might also involve in the production of pyruvate to maintain TCA cycle. Consistent with these results, we observed a marginal induction of the GABA shunt. The induction of GABA shunt may serves as an alternative source for carbon in TCA cycle and support respiratory carbon metabolism under salt stress (Che-Othman et al., 2019; Seifikalhor et al., 2019). Further, we observed marginal increase in metabolites of glycerol pathway, which are essential in providing carbon source to glycolysis and triglyceride pathways. Moreover, glycerol is known to function as intra-cellular osmoticum under salt stress (Bahieldin et al., 2013; Igamberdiev Kleczkowski, 2018).

### The correlation analysis between metabolites provides new insights for salinity studies in Pongmaia

In leaves, a strong correlation was observed among carbohydrates, fatty acids and polyols in treated plants suggesting that these metabolites may serve as first line ‘osmolyte defense’ against Na^+^ ion osmotic imbalance, protecting the structural and functional integrity of the cell and cell membrane from osmotic imbalance (Gao et al., 2013; Conde et al., 2015) (Figure 7). Further, the positive interaction between carbohydrates, intermediates of TCA cycle and polyols could serve as second line ‘ion homeostasis defense’ to maintain the intracellular pH and ion balance under salt stress (Guo et al., 2015) (Figure 7). The positive correlation, observed between carbohydrates, polyols, amino acids and organic acids might serve as third line ‘quick energy defense’ suggesting an anaplerotic role for the TCA cycle and provide immediate carbon energy source under salt stress (Zhang et al., 2017). The association between carbohydrates and amino acids showed maintain C:N balance, which may ameliorate salt-induced stress in Pongamia (Nasir et al., 2010;Naliwajski Skłodowska, 2018).

In roots, the positive correlation between organic acids, carbohydrates and fatty acids may serve as line of ‘ion homeostasis and osmolyte defense’, regulating cellular pH, osmotic potential and membrane fluidity (Sakamoto Murata, 2002; Guo et al., 2015). The enhanced correlation of secondary metabolites with these metabolites could serves as second line of ‘antioxidant and cell wall barrier defense’, where it enhances the tolerance to salinity by regulating pathways such as ROS detoxification and cell wall barrier synthesis (lignin biosynthesis) (Liu et al., 2018; Yang et al., 2018) (Figure 7). The positive association of polyols may provide a continuous osmoticum at cellular level to mitigate the osmotic imbalance raised by the increase uptake of Na^+^ ions into the vacuole (Slama et al., 2015).

### The correlation analysis between phytohormones and metabolites provides new insights for salinity studies

Hormones association with fatty acids may favour the protein lipid modifications such as S-acylation and N-myristoylation, which correspond to the irreversible link of lipid (fatty acid) to the N-terminal amino acid residue of proteins. The lipid modification could be involved in regulation of several biological processes redox homeostasis, protein-membrane associations and protein stability under various environmental stresses (Boyle et al., 2016; Majeran et al., 2018). The exogenous application of MeJA on shoots of soybean showed increased accumulation of cell wall SFAs to maintain growth under drought stress (Mohamed Latif, 2017). Further, JA also showed strong interaction with organic acids and amino acids with the salinity treatment time suggesting that organic acids and amino acids may be involved in promoting salt-induced growth by regulating osmotic balance, ion homeostasis, carbon and nitrogen balance under salt stress (Sharma et al., 2018; Siddiqi Husen, 2019). The results suggest that the increased interaction between hormones and metabolites may be beneficial for Pongamia to survive under extreme salinity stress (Sahoo et al., 2014; Formentin et al., 2018).

The number of interactions between hormones and metabolites increased with increasing treatment time in roots of 300 mM NaCl treated plants. In contrast, the number interactions were decreased with the treatment time in roots of 500 mM NaCl treated plants. The high correlation between hormones and cell-wall carbohydrates and organic acids may enhance the water permeability as well as pH regulation under salt tress conditions (Marriboina et al., 2017; Marriboina Attipalli, 2020a). The increased association of hormones with organic acids may provide immediate source of carbon energy and osmotic balance under salinity stress conditions (Assaha et al., 2017; Böhm et al., 2018; Yang Guo, 2018).

### Salinity-induced alteration in expression of ion transporter genes

In the present study, Pongamia exhibited tissue specific expression of salt-responsive transporter genes including SOS1, NHXs, PM-H^+^-ATPases, V-H^+^-ATPases, CNGCs, and other transporter genes. The expression patterns of SOS1 and SOS1-like genes correlated with Na^+^ fluoresence andNa^+^ion content,indicating that SOS1 genes are responsible for low Na^+^ levels in the leaves of Pongamia by increasing Na^+^ loading into the xylem/ apoplastic region. Further, the expression levels of SOS1 and SOS1-like genes increased significantly in salt-treated roots, which may contribute to apoplastic Na^+^ depostion under high salinity. The differential expression of SOS2 and SOS3in the both leaves and roots suggest their crucial role in salt tolerance of Pongamia through SOS pathway (Marriboina et al., 2017). In this study, we observed that the expression of NHX1 was unchaged upon 300 mM salt stress in both leaves and roots of Pongamia, while there was significant induction in 500 mM NaCl stress in both leaves and roots, which correlates with the low levels of Na^+^ fluoresence intensity and Na^+^content data in leaves of 300 mM NaCl treated plants. At high salt concentration (500 mM NaCl), Pongmia might induce NHX1 experssion to sequester Na^+^ ions rapidly into the vacuole to mitigate the Na^+^ toxicity. Similarly, in roots, induced NHX1 expression may indicate the vacuolar Na^+^ sequestration. In contrast, NHX1 levels wereunaffected at intial impostion of stress, suggesting involvement of other NHXs isofoms for vacuolar Na^+^ sequestration at intial stages of salt stress. The differential expresssion of other NHXs isoforms such as NHX3, NHX6 nd NHX6-like in both leaves and roots suggests their possible roles in ion homeostasis, plant growth and development under salt stress (Bassil et al., 2018; Dragwidge et al., 2018).

In this study,expression of HKT1:1 followed similar trend in both leaves and roots of salt stressed plants. Previous studies suggested that HKT family transporter proteins can mediate exclusion of Na^+^from leaves and roots by translocating shoot-to-rootand root-to-shoot Na^+^ delivery by withdrwaing Na^+^ from xylem stream into phloem (Munns et al., 2012; Hill et al., 2013; An et al., 2017; Zhang et al., 2018). Induction of HKT1:1 expression upon initial imposition of salt stress in both leaves and roots may promote Na^+^ exclusion tolimit Na^+^ toxcity in the respective tissues(Zhang et al., 2018). However, with increasing salt stress treatment time, the marginal expression of HKT1:1genemay regulate the retrevial of Na^+^ from the xylem (Davenport et al., 2007; Ali et al., 2016). The expression levels of CLC1 indicates that CLC1 may not be involved in the Cl^-^ vacuolar sequestation (Wei et al., 2016).

PM-H^+^-ATPase family pumps playa crucial role in improving the salt tolerance by maintaing the intracellular pH balance, transmembrane potential and ion homeostasis under salt stress conditions (Olfatmiri et al., 2014;Falhof et al., 2016; Shabala et al., 2016). The diffrences in the expression levels of PM-H^+^-ATPase isoforms could be due toconfigurational and/or post-translational modifications of these isforms under salt stress, which could promote growth under salinity stress. The increased expression of V-H^+^-PPase and V-H^+^-ATPaseB and E subunit in leaves of salt treated plants may control the depolarization of vacuolar membrane potential, which is generated by excess depostion of Na^+^ ions in leaves of Pongamia upon prolonged salt exposure (Graus et al., 2018; Marriboina Attipalli, 2020a).

An increased expression levels of V-CHX1 may involve in plant growth by improving cellular ion homeostasis, pH balance and osmoregulation under saline conditions (Guan et al., 2014; Qi et al., 2014; Liu et al., 2017). Enhanced experssion of CCX1 may involved in regulation of intracellular Ca^2+^ levels, which may help in vacuolar Na^+^ sequestration, ROS and ion homeostasis under salt stress (Yong et al., 2014; Li et al., 2016; Corso et al., 2018). Differential expression of CNGC5, CNGC17 and CNGC17-like may induce Ca^2+^ derived reponses to mitigate the negative effects of salt stress (Wang et al., 2013; Saand et al., 2015).

### Salinity-induced alteration in expression of antioxidant genes

Excess Na^+^ accumulation in the plant causes ionic imbalance which results in ion toxicity, oxidative stress, and generation reactive oxygen species (ROS) (AbdElgawad et al., 2016). To combat the ROS induced cellular damage plant induces several antioxidant enzymes such as superoxide dismutase (SOD), catalase (CAT) and peroxidase (POD) (Reddy et al., 2004). In this study, the observed higher expression levels of the different SOD isoforms might protect the plants at the onset of oxidative stress much before the cellular antioxidant signaling cascade initiates its function under high salinity. According to Gill Tuteja, (2010) the H_2_O_2_ produced through SOD activity, provides an additional advantage to the plant by strengthening cell wall through lignin biosynthesis (Figure 7). Also, higher CAT4 expression levels in roots might help the plant to reduce the higher levels of H_2_O_2_ under high salt conditions. The lower expression levels of CAT4 in the leaves may reflect the dynamic and tissue specificity of the antioxidant system under high salt environment (AlHassan et al., 2017). Increased expression of POD in leaves and roots were well correlated with the respective lignin depositions. The classIII apoplastic peroxidases play a major role in maintaining the structural integrity of the cell wall under salt stress (Kim et al., 2012). In addition, the expression levels of several other cell-wall modifying and lignin biosynthesis genes were monitored which include, PAL1, PAL2, CAD1, CAD1-like, CAD2, and CAD6. The increased expression levels of both PAL1 and PAL2 in 500 mM NaCl treated roots might permit the plant to preserve the cell wall integrity under high salt environment (Lee et al., 2007). The consistent increase in the CAD1-like gene expression suggests its possible role in reinforcing the strength and rigidity of the cell wall to enhance the selectivity and permeability of cell wall for solute and water transport under high saline environment (Zhao et al., 2013).

## Conclusions

This is the first study explicitly focusing on salt tolerance mechanisms in *Pongamia pinnata*.. Salinity induced Ca^2+^ levels not only counteract the Na^+^ toxicity but also activate numerous signaling pathways including JAs signaling. Based on our correlation results, we hypothesize that three phytohormones JAs, SA and ABA play crucial role in the adaptation of Pongamia plants to high salinity stress. On the other hand, induction of metabolites such as a sugars, amino acids and polyols provides constant supply of nutrients and are also involved in conferring osmotic and ionic homeostasis under salinity stress. The downstream effect of the above signaling pathways could influence the defense (antioxidant) and transporter gene expression patterns conferring redox homeostasis under salinity stress, ultimately leading to Na^+^ ion exclusion and sequestration in the apoplasmic regions.

## Author contribution

S.M. designed the research; S.M. and K.S. performed the research; A.D.Y., K.S. and S.M. performed bioinformatics analyses; S.M. and D.S. analysed the data; S.M., and D.S. wrote the paper; A.R.R. supervised the research and R.P.S. helped in the discussion.

## Acknowledgments

We gratefully acknowledge Tree Oils India Limited (TOIL), Zaheerabad, Telangana State, India for providing Pongamia seeds. S.M. acknowledges University Grant Commission (UGC), New Delhi, India for fellowship.

## Supplementary Data

**Figure S1.** Effect of salt stress on plant morphology

**Figure S2.** Effect of salt stress on photosynthetic performance, chlorophyll fluorescence, chlorophyll content and ion contents of (A) Responses of carbon exchange rate (CER) (µmol m^-2^ s^-1^), (B) changes in maximal photochemical efficiency of PSII (Fv/Fm), (C) total chlorophyll content, (D) relative water content in the leaves and roots of control and salt-treated plants (RWC) (%), (E) Cl^-^ ion content in the leaves and roots of control and salt-treated plants, (F) Ca^2+^ ion content in the leaves and roots of control and salt-treated plants, (G) Na^+^ in the leaves and roots of control and salt-treated plants, (H) and (I) variations in the ratios of Na^+^/Cl^-^ and Na^+^/Ca^2+^ ratios in the leaves and roots of control and salt-treated plants. Error bars represent the mean ± SD (n=6). Two-way ANOVA test was performed to measure P-values ns (not significant) * (P<0.05), ** (P<0.01) and * * * (p<0.001) respectively.

**Figure S3.** Illumination of Na^+^ ion fluorescence in leaves of control and salt-treated *P. pinnata*. Confocal images of leaves were stained with CoroNa-Green AM (green colour) and propidium iodide (red colour). (A, B and C) cross sections of leaves of control plants, (D, E and F), cross sections of leaves of 300 mM NaCl 1, 4 and 8DAS, (G, H and I) cross sections of leaves 500 mM NaCl 1, 4 and 8DAS respectively. Red arrows were drawn to measure Na^+^ fluorescence intensity. Quantification of Na^+^ fluorescence intensity more than 12 images were pooled from five biological replicates in control and salt-treated plants. Error bar represents the mean ± SD (n=12). Two-way ANOVA test was performed to measure P-values ns (not significant) * (P<0.05), ** (P<0.01) and * * * (p<0.001) respectively.

**Figure S4.** Illumination of Na^+^ ion fluorescence in roots of control and salt-treated *P. pinnata*. Confocal images of roots were stained with CoroNa-Green AM (green colour) and propidium iodide (red colour). (A, B and C) cross sections of roots of control plants, (D, E and F), cross sections of roots of 300 mM NaCl 1, 4 and 8DAS, (G, H and I) cross sections of roots 500 mM NaCl 1, 4 and 8DAS, (J, K and L) magnified images of roots of control plants, (M, N and O) magnified images of roots of 300 mM NaCl 1, 4 and 8DAS, (P, Q and R) magnified images of roots 500 mM NaCl 1, 4 and 8DAS respectively. White arrows were drawn to measure Na^+^ fluorescence intensity. Quantification of Na^+^ fluorescence intensity more than 12 images were pooled from five biological replicates in control and salt-treated plants. Error bar represents the mean ± SD (n=12). Two-way ANOVA test was performed to measure P-values ns (not significant) * (P<0.05), ** (P<0.01) and * * * (p<0.001) respectively.

**Figure S5.** Heat map of metabolite changes in leaves of *P. pinnata* under salt stress. Log_2_ ratios of the relative fold change values (1 to 12) of each metabolite is represented in the form of a single horizontal row portioned with six columns of two salt treatments (300 and 500 mM NaCl) and three different time points (1, 4 and 8DAS) respectively. Each metabolite value is an average of six biological replicates represented with different colour codes red or blue according to the scale bar. The colour scale indicates degree of correlation.

**Figure S6.** Heat map of metabolite changes in roots of *P. pinnata* under salt stress. Log_2_ ratios of the relative fold change values (1 to 12) of each metabolite is represented in the form of a single horizontal row portioned with six columns of two salt treatments (300 and 500 mM NaCl) and three different time points (1, 4 and 8DAS) respectively. Each metabolite value is an average of six biological replicates represented with different colour codes red or blue according to the scale bar. The colour scale indicates degree of correlation.

**Figure S7.** Representative figure of semi-quantitative PCR analysis of transporter genes expression in leaves and roots of *P. pinnata* under three different salinity stress conditions 0, 300 and 500 mM NaCl at 1, 4 and 8DAS. The expression of levels of each gene normalized by using reference gene 18s rRNA.

**Figure S8.** The band intensities of transporter genes of semi-quantitative PCR analysis (showed in **Supplementary Figure 7)** in leaves and roots of *P. pinnata* under three different salinity stress conditions 0, 300 and 500 mM NaCl at 1, 4 and 8DAS respectively.

**Figure S9**. Relative mRNA expression levels of cell-wall enzymes and antioxidant enzymes in leaves of *P. pinnata* under Salt Stress Conditions. Log_2_ fold changes of CAT4, FeSOD, MnSOD, Cu/ZnSOD1, Cu/ZnSOD1-like, POD, PAL1, PAL2, CAD2, and CAD6, in leaves (A) and roots (B) of salt-treated plants of *P. pinnata* at 1, 4 and 8DAS respectively when compared to their corresponding controls. Error bar represents the mean ± SD (n=6).

**Figure S10.** Representative figure of semi-quantitative PCR analysis of cell-wall enzymes and antioxidant enzyme genes expression in leaves and roots of *P. pinnata* under three different salinity stress conditions 0, 300 and 500 mM NaCl at 1, 4 and 8DAS. The expression of levels of each gene normalized by using reference gene 18s rRNA.

**Figure S11.** The band intensities of CAD1 and CAD1-like genes in leaves and roots of *P. pinnata* under three different salinity stress conditions 0, 300 and 500 mM NaCl at 1, 4 and 8DAS respectively.

**Table S1.** Relative concentration and fold changes of major metabolites in leaves of *P. pinnata* under three different salinity stress conditions 0 (Control), 300 and 500 mM NaCl at 1, 4 and 8DAS. Six biological replicates were used to measure the relative fold changes of each metabolite and two-way ANOVA test was performed to measure P-values ns (not significant) * (P<0.05), ** (P<0.01) and * * * (p<0.001) respectively.

**Table S2.** Relative concentration and fold changes of major metabolites in roots of *P. pinnata* under three different salinity stress conditions 0 (Control), 300 and 500 mM NaCl at 1, 4 and 8DAS. Six biological replicates were used to measure the relative fold changes of each metabolite and two-way ANOVA test was performed to measure P-values ns (not significant) * (P<0.05), ** (P<0.01) and * * * (p<0.001) respectively.

**Table S3.** List of all 71 metabolites of 300 mM NaCl treated leaves of *P. pinnata* at 1, 4 and 8DAS and their abbreviations showed in figure 3 A-F.

**Table S4.** List of all 71 metabolites of 500 mM NaCl treated leaves of *P. pinnata* at 1, 4 and 8DAS and their abbreviations showed in figure 3 G-L.

**Table S5.** List of all 71 metabolites of 300 mM NaCl treated roots of *P. pinnata* at 1, 4 and 8DAS and their abbreviations showed in figure 4 A-F.

**Table S6.** List of all 71 metabolites of 500 mM NaCl treated roots of *P. pinnata* at 1, 4 and 8DAS and their abbreviations showed in figure 4 G-L.

**Table S7**. List of primers were used in semi-quantitative-PCR, RT-PCR analysis to measure mRNA expression.

## References

Abdallaha MMS, Abdelgawad ZA, El-Bassiounya HMS. 2016. Alleviation of the adverse effects of salinity stress using trehalose in two rice varieties. South African Journal of Botany 103, 275–282.

AbdElgawad H, Zinta G, Hegab MM, Pandey R, Asard H, Abuelsoud W. 2016. High salinity induces different oxidative stress and antioxidant responses in maize plants organs. Frontiers in Plant Science 7, 276.

Abebe T, Guenzi AC, Martin B, Cushman JC. 2003. Tolerance of mannitol accumulating transgenic wheat to water stress and salinity. Plant Physiology 131, 1748–1755.

AlHassan M, Chaura J, Donat-Torres MP, Boscaiu M, Vicente O. 2017. Antioxidant responses under salinity and drought in three closely related wild monocots with different ecological optima. AoB Plants 9, plx009.

Ali A, Raddatz N, Aman R, Kim S, Park HC, Jan M, Bressan RA. 2016. A single amino acid substitution in the sodium transporter HKT1 associated with plant salt tolerance. Plant Physiology 171, 2112–2126.

Ali MF, Baek KH. 2020. Jasmonic acid signaling pathway in response to abiotic stresses in plants. International Journal of Molecular Sciences 21, 621.

An D, Chen JG, Gao YQ, Li X, Chao ZF, Chen ZR, Li QQ, Han ML, Wang YL, Wang YF, Chao DY. 2017. *AtHKT1* drives adaptation of *Arabidopsis thaliana* to salinity by reducing floral sodium content. PLoS Genetics 13, e1007086.

Anower MR, Peel MD, Mott IW, Wu Y. 2017. Physiological process associated with salinity tolerance in an alfalfa half-sib family. Journal of Agronomy and Crop Science 203, 506–518.

Arnon DI. 1949. Copper enzymes in isolated chloroplasts polyphenol oxidase in *Beta vulgaris*. Plant Physiology 24, 1–15.

Assaha DV, Ueda A, Saneoka H, Al□Yahyai R, Yaish MW. 2017. The role of Na^+^ and K^+^ transporters in salt stress adaptation in glycophytes. Frontiers in Physiology 8, 509.

Atikij T, Syaputri Y, Iwahashi H, Praneenararat T, Sirisattha S, Kageyama H, Sirisattha, RW. 2019. Enhanced lipid production and molecular dynamics under salinity stress in green microalga *Chlamydomonas reinhardtii* (137C). Marine Drugs 17, 484.

Baetz U, Eisenach C, Tohge T, Martinoia E, De-Angeli A. 2016. Vacuolar chloride fluxes impact ion content and distribution during early salinity stress. Plant Physiology 172, 1167–1181.

Bahieldin A, Sabir JSM, Ramadan A, Alzohairy AM, Younis RA, Shokry AM, Gadalla NO, Edris S, Hassan SM, Al-Kordy MA, Kamal KBH, Rabah S, Abuzinadah OA, El-Domyati FM. 2013. Control of glycerol biosynthesis under high salt stress in *Arabidopsis*. Functional Plant Biology 41, 87.

Bassil E, Zhang S, Gong H, Tajima H, Blumwald E. 2018. Cation specificity of vacuolar NHX-type cation/H^+^ antiporters. Plant Physiology 179, 616–629.

Böhm J, Messerer M, Müller HM, Scholz-Starke J, Gradogna A, Scherzer S, Maierhofer T, Bazihizina N, Zhang H, Stigloher C, Ache P, Al-Rasheid KAS, Mayer KFX, Shabala S, Carpaneto A, Haberer G, Zhu JK, Hedrich R. 2018. Understanding the molecular basis of salt sequestration in epidermal bladder cells of *Chenopodium quinoa*. Current Biology 28, 3075–3085.

Boyle PC, Schwizer S, Hind SR, Kraus CM, De-la-Torre-Diaz S, He B, Martin GB. 2016. Detecting N-myristoylation and S-acylation of host and pathogen proteins in plants using click chemistry. Plant Methods 12, 38.

Cai Q, Yuan Z, Chen M, Yin C, Luo Z, Zhao X, Liang W, Hu J, Zhang D. 2014. Jasmonic acid regulates spikelet development in rice. Nature Communications 5, 3476.

Cao D, Lutz A, Hill CB, Callahan DL, Roessner U. 2017. A quantitative profiling method of phytohormones and other metabolites applied to barley roots subjected to salinity stress. Frontiers in Plant Science 7, 2070.

Cao J, Li M, Chen J, Liu P, Li Z. 2016. Effects of MeJA on *Arabidopsis* metabolome under endogenous JA deficiency. Scientific Reports 6, 37674.

Chen X, Wang H, Li X, Ma K, Zhan Y, Zeng F. 2019. Molecular cloning and functional analysis of 4-coumarate:CoA ligase 4 (4CL-like 1) from *Fraxinus mandshurica* and its role in abiotic stress tolerance and cell wall synthesis. BMC Plant Biology 19, 231.

Che-Othman MH, Jacoby RP, Millar AH, Taylor NL. 2019. Wheat mitochondrial respiration shifts from the TCA cycle to the GABA shunt under salt stress. New Phytologist 225, 1166–1180.

Choi WG, Toyota M, Kim SH, Hilleary R, Gilroy S. 2014. Salt stress-induced Ca^2+^ waves are associated with rapid, long-distance root-to-shoot signalling in plants. Proceedings of the National Academy of Sciences of the United States of America 111, 6497–6502.

Conde A, Regalado A, Rodrigues D, Costa JM, Blumwald E, Chaves MM. 2015. Polyols in grape berry: transport and metabolic adjustments as a physiological strategy for water-deficit stress tolerance in grapevine. Journal of Experimental Botany 66, 889–906.

Corso M, Doccula FG, de-Melo JRF, Costa A, Verbruggen N. 2018. Endoplasmic reticulum-localized CCX2 is required for osmotolerance by regulating ER and cytosolic Ca^2+^ dynamics in *Arabidopsis*. Proceedings of the National Academy of Sciences of the United States of America 115, 3966–3971.

Davenport RJ, Muñoz-Mayor A, Jha D, Essah PA, Rus A, Tester M. 2007. The Na^+^ transporter AtHKT1;1 controls retrieval of Na^+^ from the xylem in *Arabidopsis*. Plant Cell Environment 30, 497–507.

Devinar G, Llanes A, Masciarelli O, Luna V. 2013. Different relative humidity conditions combined with chloride and sulfate salinity treatments modify abscisic acid and salicylic acid levels in the halophyte *Prosopis strombulifera*. Plant Growth Regulation 70, 247–256.

Diallo AO, Agharbaoui Z, Badawi MA, Ali-Benali MA, Moheb A, Houde M, Sarhan F. 2014. Transcriptome analysis of an *mvp* mutant reveals important changes in global gene expression and a role for methyl jasmonate in vernalization and flowering in wheat. Journal of Experimental Botany 65, 2271–2286.

Dragwidge JM, Ford BA, Ashnest JR, Das P, Gendall AR. 2018. Two endosomal NHX-type Na^+^/H^+^ antiporters are involved in auxin mediated development in *Arabidopsis thaliana*. Plant Cell Physiology 59, 1660–1669.

Dumschott K, Dechorgnat J, Merchant A. 2019. Water deficit elicits a transcriptional response of genes governing D-pinitol biosynthesis in soybean (*Glycine max*). International Journal of Molecular Sciences 20, 2411.

Fahad S, Nie L, Chen Y, Wu C, Xiong D, Saud S, Hongyan L, Cui K, Huang J. 2015. Crop plant hormones and environmental stress. Sustainable Agriculture Reviews 15, 371–400.

Falhof J, Pedersen JT, Fuglsang AT, Palmgren M. 2016. Plasma membrane H^+^-ATPase regulation in the center of plant physiology. Molecular Plant 9, 323–337.

Fattorini L, Hause B, Gutierrez L, Veloccia A, Rovere FD, Piacentini D, Falasca G, Altamura MM. 2018. Jasmonate promotes auxin-induced adventitious rooting in dark-grown *Arabidopsis thaliana* plants and stem thin cell layers by a cross-talk with ethylene signaling and a modulation of xylogenesis. BMC Plant Biology 18, 182.

Felle HH, Hermann A, Hückelhoven R, Kogel KH. 2005. Root-to-shoot signalling: apoplastic alkalinization, a general stress response and defence factor in barley (*Hordeum vulgare*). Protoplasma 227, 17–24.

Formentin E, Barizza E, Stevanato P, Falda M, Massa F, Tarkowskà D, Novak O, Lo-Schiavo F. 2018. Fast regulation of hormone metabolism contributes to salt tolerance in rice (*Oryzasativa* spp. Japonica, L.) by inducing specific morpho-physiological responses. Plants (Basel) 7, 75.

Fukuda A, Tanaka Y. 2006. Effects of ABA, auxin, and gibberellin on the expression of genes for vacuolar H^+^-inorganic pyrophosphatase, H^+^-ATPase subunit A, and Na^+^/H^+^ antiporter in barley. Plant Physiology and Biochemistry 44, 351–358.

Gao R, Duan K, Guo G, Du Z, Chen Z, Li L, He T, Lu R, Huang J. 2013. Comparative transcriptional profiling of two contrasting barley genotypes under salinity stress during the seedling stage. International Journal of Genomics 139, 822–835.

Geilfus CM, Tenhaken R, Carpentier SC. 2017. Transient alkalinization of the leaf apoplast stiffens the cell wall during onset of chloride-salinity in corn leaves. Journal of Biological Chemistry 292, 18800–18813.

Ghanem ME, Albacete A, Smigocki AC, Frébort I, Pospíšilová H, Martínez□Andújar C, Acosta M, Sánchez-Bravo J, Lutts S, Dodd IC, Pérez-Alfocea F. 2011. Root[synthesized cytokinins improve shoot growth and fruit yield in salinized tomato (*Solanum lycopersicum* L.) plants. Journal of Experimental Botany 62, 125–140.

Gharbi E, Lutts S, Dailly H, Quinet M. 2018. Comparison between the impacts of two different modes of salicylic acid application on tomato (*Solanum lycopersicum*) responses to salinity. Plant Signaling Behavior 13, e146936.

Gilbert L, Alhagdow M, Nunes-Nesi A, Quemener B, Guillon F, Bouchet B, Faurobert M, Gouble B, Page D, Garcia V, Petit J, Stevens R, Causse M, Fernie AR, Lahaye M, Rothan C, Baldet P. 2009. GDP-D-mannose 3, 5-epimerase (GME) plays a key role at the intersection of ascorbate and non-cellulosic cell-wall biosynthesis in tomato. The Plant Journal 60, 499–508.

Gill SS, Tuteja N. 2010. Reactive oxygen species and antioxidant machinery in abiotic stress tolerance in crop plants. Plant Physiology and Biochemistry 48, 909–930.

Golan Y, Shirron N, Avni A, Shmoish M, Gepstein S. 2016. Cytokinins induce transcriptional reprograming and improve Arabidopsis plant performance under drought and salt stress conditions. Frontiers in Environmental Science 4, 63.

Gonzalez P, Syvertsen JP, Etxeberria E. 2012. Sodium distribution in salt-stressed citrus root stock plants. Horticultural Science 47, 1504–1511.

Graus D, Konrad KR, Bemm F, Patir-Nebioglu MG, Lorey C, Duscha K, Güthoff T, Herrmann J, Ferjani A, Cuin TA, Roelfsema MRG, Schumacher K, Neuhaus HE, Marten I, Hedrich R. 2018. High V-PPase activity is beneficial under high salt loads, but detrimental without salinity. New Phytologist 219, 1421–1432.

Guan R, Qu Y, Guo Y, Yu L, Liu Y, Jiang J, Chen J, Ren Y, Liu G, Tian L, Jin L, Liu Z, Hong H, Chang R, Gilliham M, Qiu L. 2014. Salinity tolerance in soybean is modulated by natural variation in *GmSALT3*. Plant Journal 80, 937–950.

Guo M, Lu JP, Zhai YF, Chai WG, Gong ZH, Lu MH. 2015. Genome-wide analysis, expression profile of heat shock factor gene family (CaHsfs) and characterization of CaHsfA2 in pepper (*Capsicum annuum* L.). BMC Plant Biology 15, 151.

Gupta A, Hisano H, Hojo Y, Matsuura T, Ikeda Y, Mori IC, Senthil-Kumar M. 2017. Global profiling of phytohormone dynamics during combined drought and pathogen stress in Arabidopsis thaliana reveals ABA and JA as major regulators. Scientific Reports 7, 4017.

Hanin M, Ebel C, Ngom M, Laplaze L, Masmoudi K. 2016. New insights on plant salt tolerance mechanisms and their potential use for breeding. Frontiers in Plant Science 7, 1787.

Hill CB, Jha D, Bacic A, Tester M, Roessner U. 2013. Characterization of ion contents and metabolic responses to salt stress of different *Arabidopsis* AtHKT1;1 genotypes and their parental strains. Molecular Plant 6, 350–368.

Hossain MS, Persicke M, ElSayed AI, Kalinowski J, Dietz KJ. 2017. Metabolite profiling at the cellular and subcellular level reveals metabolites associated with salinity tolerance in carbohydrate beet. Journal of Experimental Botany 68, 5961–5976.

Huang J, Lu X, Yan H, Chen S, Zhang W, Huang R, Zheng Y. 2012. Transcriptome characterization and sequencing-based identification of salt responsive genes in *Millettia pinnata*, a semi-mangrove plant. DNA Research 19, 195–207.

Husen A, Iqbal M, Sohrab SS, Ansari MKA. 2018. Salicylic acid alleviates salinity caused damage to foliar functions, plant growth and antioxidant system in Ethiopian mustard (*Brassica carinataa*. Br.). Agriculture Food Security 7, 44.

Igamberdiev AU, Kleczkowski LA. 2018. The glycerate and phosphorylated pathways of serine synthesis in plants: the branches of plant glycolysis linking carbon and nitrogen metabolism. Frontiers in Plant Science 9, 318.

Ishimaru Y, Hayashi K, Suzuki T, Fukaki H, Prusinska J, Meester C, Quareshy M, Egoshi S, Matsuura H, Takahashi K, Kato N, Kombrink E, Napier RM, Hayashi KI, Uedaa M. 2018. Jasmonic acid inhibits auxin-induced lateral rooting independently of the CORONATINE INSENSITIVE1 receptor. Plant Physiology 177, 1704–1716.

Jang G, Chang SH, Um TY, Lee S, Kim JK, Do-Choi Y. 2017. Antagonistic interaction between jasmonic acid and cytokinin in xylem development. Scientific Reports 7, 10212.

Jayakannan M, Bose J, Babourina O, Rengel Z, Shabala S. 2013. Salicylic acid improves salinity tolerance in *Arabidopsis* by restoring membrane potential and preventing salt-induced K^+^ loss via a GORK channel. Journal of Experimental Botany 64, 2255–2268.

Kang DJ, Seo YJ, Lee JD, Ishii R, Kim KU, Shin DH, Park SK, Jang SW, Lee IJ. 2005. Jasmonic acid differentially affects growth, ion uptake and abscisic acid concentration in salt-tolerant and salt-sensitive rice cultivars. Journal of Agronomy Crop Science 191, 273–282.

Kim BH, Kim SY, Nam KH. 2012. Genes encoding plant-specific class III peroxidases are responsible for increased cold tolerance of the brassinosteroid-insensitive 1 mutant. Molecules Cells 34, 539–548.

Kim JI, Baek D, Park HC, Chun HJ, Oh DH, Lee MK, Cha JY, Kim WY, Kim MC, Chung WS, Bohnert HJ, Lee SY, Bressan RA, Lee SW, Yun DJ. 2012. Overexpression of *Arabidopsis YUCCA6* in potato results in high-auxin developmental phenotypes and enhanced resistance to water deficit. Molecular Plant 6, 337–349.

Korkmaz D. 2001. Precipitation titration: “determination of chloride by the Mohr method”. Methods 2, 4.

Lee BR, Kim KY, Jung WJ, Avice JC, Ourry A, Kim TH. 2007. Peroxidases and lignification in relation to the intensity of water-deficit stress in white clover (*Trifoliumrepens* L.). Journal of Experimental Botany 58, 1271–1271.

Li Q, Zheng J, Li S, Huang G, Skilling SJ, Wang L, Li L, Li M, Yuan L, Liu P. 2017. Transporter-mediated nuclear entry of jasmonoyl-isoleucine is essential for jasmonate signalling. Molecular Plant 10, 695–708.

Li Z, Wang X, Chen J, Gao J, Zhou X, Kuai B. 2016. CCX1, a putative Cation/Ca^2+^ exchanger, participates in regulation of reactive oxygen species homeostasis and leaf senescence. Plant Cell Physiology 57, 2611–2619.

Liu J, Gao F, Ren J, Lu X, Ren G, Wang R. 2017. A novel AP2/ERF transcription factor CR1 regulates the accumulation of vindoline and serpentine in *Catharanthus roseus*. Frontiers in Plant Science 8, 2082.

Liu Q, Luo L, Zheng L. 2018. Lignins: biosynthesis and biological functions in plants. International Journal of Molecular Sciences 19, 335.

Liu YD, Yin ZJ, Yu JW, Li J, Wei HL, Han XL, Shen FF. 2012. Improved salt tolerance and delayed leaf senescence in transgenic cotton expressing the *Agrobacterium IPT* gene. Biologia Plantarum 56, 237–246.

Livak KJ, Schmittgen TD. 2001. Analysis of relative gene expression data using real-time quantitative PCR and the 2-ΔΔCT method. Methods 25, 402–408.

Lombardo VA, Osorio S, Borsani J, Lauxmann MA, Bustamante CA, Budde CO, Andreo CS, Lara MV, Fernie AR, Drincovich MF. 2011. Metabolic profiling during peach fruit development and ripening reveals the metabolic networks that underpin each developmental stage. Plant Physiology 157, 1969–1710.

Majeran W, Le-Caer JP, Ponnala L, Meinnel T, Giglione C. 2018. Targeted profiling of *Arabidopsis thaliana* sub-proteomes illuminates co- and post-translationally N-Terminal myristoylated proteins. The Plant Cell 30, 543–562.

Marriboina S, Attipalli RR. 2020a. Hydrophobic cell-wall barriers and vacuolar sequestration of Na^+^ ions are the key mechanisms conferring high salinity tolerance in a biofuel tree species *Pongamia pinnata* L. pierre. Environmental Experimental Botany 171, 103949.

Marriboina S, Attipalli RR. 2020b. Optimization of hydroponic growth system and Na^+^-fluorescence measurements for tree species *Pongamia pinnata* (L.) pierre. MethodsX 7, 100809.

Marriboina S, Sengupta D, Kumar S, Attipalli RR. 2017. Physiological and molecular insights into the high salinity tolerance of *Pongamia pinnata* (L.) pierre, a potential biofuel tree species. Plant Science 258, 102–111.

Maury S, Sow MD, Le-Gac AL, Genitoni J, Lafon-Placette C, Mozgova I. 2019. Phytohormone and chromatin crosstalk: The missing link for developmental plasticity?. Frontiers in Plant Science 10, 395.

Md-Hossain S. 2019. Present scenario of global salt affected soils, its management and importance of salinity research. International Research Journal of Biological Sciences 1, 1–3.

Melo YL, Dantas CVS, Melo YL, Maia JM, De-Macêdo CEC. 2016. Changes in osmotic and ionic indicators in *Ananascomosus* (L.) cv. MD gold pre-treated with phytohormones and submitted to saline medium. The Revista Brasileira de Fruticultura 39, e–155.

Mitra S, Baldwin IT. 2014. RuBPCase activase (RCA) mediates growth-defense trade-offs: silencing RCA redirects jasmonic acid (JA) flux from JA-isoleucine to methyl jasmonate (MeJA) to attenuate induced defense responses in *Nicotiana attenuata*. New Phytologist 201, 1385–1395.

Mohamed IH, Latif HH. 2017. Improvement of drought tolerance of soybean plants by using methyl jasmonate. Physiology and Molecular Biology of Plants 23, 545–556.

Moing A, Aharoni A, Biais B, Rogachev I, Meir S, Brodsky L, Allwood JW, Erban A, Dunn WB, Kay L, de-Koning S, de-Vos RC, Jonker H, Mumm R, Deborde C, Maucourt M, Bernillon S, Gibon Y, Hansen TH, Husted S, Goodacre R, Kopka J, Schjoerring JK, Rolin D, Hall RD. 2011. Extensive metabolic cross-talk in melon fruit revealed by spatial and developmental combinatorial metabolomics. New Phytologist 190, 683–696.

Morton MJL, Awlia M, Al-Tamimi N, Saade S, Pailles Y, Negrão S, Tester M. 2019. Salt stress under the scalpel-dissecting the genetics of salt tolerance. The Plant Journal 97, 148–163.

Munemasa S, Hossain MA, Nakamura Y, Mori IC, Murata Y. 2011. The *Arabidopsis* calcium-dependent protein kinase, CPK6, functions as a positive regulator of methyl jasmonate signaling in guard cells. Plant Physiology 155, 553–561.

Munns R, James RA, Xu B, Athman A, Conn SJ, Jordans C, Byrt CS, Hare RA, Tyerman SD, Tester M, Plett D, Gilliham M. 2012. Wheat grain yield on saline soils is improved by an ancestral Na^+^ transporter gene. Nature Biotechnology 30, 360–364.

Munns R, Wallace PA, Teakle NL, Colmer TD. 2010. Measuring soluble ion concentrations (Na^+^, K^+^, Cl^-^) in salt-treated plants, R. Sunkar, eds. Plant stress tolerance. Methods in molecular biology. (Springer protocols, Berlin: Humana Press), pp. 371–382.

Naliwajski MR, Skłodowska M. 2018. The relationship between carbon and nitrogen metabolism in cucumber leaves acclimated to salt stress. Peer J 6, e6043.

Nasir FA, Batarseh M, Abdel-Ghani AH, Jiries A. 2010. Free amino acids content in some halophytes under salinity stress in arid environment. Jordan. Clean Soil Air Water 38, 592–600.

Nishiyama R, Watanabe Y, Fujita Y, Le DT, Kojima M, Werner T, Vankova R, Yamaguchi-Shinozaki K, Shinozaki K, Kakimoto T, Sakakibara H, Schmülling T, Tran LSP. 2011. Analysis of cytokinin mutants and regulation of cytokinin metabolic genes reveals important regulatory roles of cytokinins in drought, salt and abscisic acid responses, and abscisic acid biosynthesis. The Plant Cell 23, 2169–2183.

Nitschke S, Cortleven A, Iven T, Feussner I, Havaux M, Riefler M, Schmülling T. 2016. Circadian stress regimes affect the circadian clock and cause jasmonic acid-dependent cell death in cytokinin-deficient *Arabidopsis* plants. The Plant Cell 28, 1616–1639.

Olfatmiri H, Alemzadeh A, Zakipour Z. 2014. Up-regulation of plasma membrane H^+^-ATPase under salt stress may enable *Aeluropus littoralis* to cope with stress. Molecular Biology Research Communications 3, 67–75.

Pan X, Welti R, Wang X. 2010. Quantitative analysis of major plant hormones in crude plant extracts by high-performance liquid chromatography-mass spectrometry. Nature Protocols 5, 986–992.

Peng Z, He SP, Sun JL, Pan ZE, Gong WF, Lu YL, De XM. 2016. Na^+^ compartmentalization related to salinity stress tolerance in upland cotton (*Gossypium hirsutum*) plants. Scientific Reports 6, 34548.

Qi X, Li MW, Xie M, Liu X, Ni M, Shao G, Song C, Yim AKY, Tao Y, Wong FL, Isobe S, Wong CF, Wong KS, Xu C, Li C, Wang Y, Guan R, Sun F, Fan G, Xiao Z, Zhou F, Phang TH, Liu X, Tong SW, Chan TF, Yiu SM, Tabata S, Wang J, Xu X, Lam HM. 2014. Identification of a novel salt tolerance gene in wild soybean by whole-genome sequencing. Nature Communications 5, 4340.

Quinn LD, Straker KC, Guo J, Kim S, Santanu T, Kling G, Lee DK, Voigt TB. 2015. Stress-tolerant feed stocks for sustainable bioenergy production on marginal land. Bioenergy Research 8, 1081–1100.

Rahneshan Z, Nasibi F, Moghadam AA. 2018. Effect of salinity stress on some growth, physiological, biochemical parameters and nutrients in two pistachio (*Pistacia vera* L.) rootstocks. Plant Environment Interactions 13, 73–82.

Raza A, Razzaq A, Mehmood SS, Zou X, Zhang X, Lv Y, Xu J. 2019. Impact of climate change on crops adaptation and strategies to tackle its outcome: A review. Plants 8, 39.

Reddy AR, Chaitanya KV, Vivekanandan M. 2004. Drought induced responses of photosynthesis and antioxidant metabolism in higher plants. Journal of Plant Physiology 161, 1189–1202.

Roessner U, Wagner C, Kopka J, Trethewey RN, Willmitzer L. 2000. Simultaneous analysis of metabolites in potato tuber by gas chromatography-mass spectrometry. The Plant Journal 23, 131–142.

Saand MA, Xu YP, Munyampundu JP, Li W, Zhang XR, Cai XZ. 2015. Phylogeny and evolution of plant cyclic nucleotide-gated ion channel (CNGC) gene family and functional analyses of tomato CNGCs. DNA Research 22, 471–483.

Sahoo RK, Ansari MW, Tuteja R, Tuteja N. 2014. *OsSUV3* transgenic rice maintains higher endogenous levels of plant hormones that mitigates adverse effects of salinity and sustains crop productivity. Rice 7, 17.

Sakamoto T, Murata N. 2002. Regulation of the desaturation of fatty acids and its role in tolerance to cold and salt stress. Current Opinion in Microbiology 5, 208–210.

Saleh L, Plieth C. 2013. *A9C* sensitive Cl^-^ accumulation in *A. thaliana* root cells during salt stress is controlled by internal and external calcium. Plant Signaling Behavior 8, e24259.

Samuel S, Scott PT, Gresshoff PM. 2013. Nodulation in the legume biofuel feedstock tree *Pongamia pinnata*. Agricultural Research 2, 207–214.

Sánchez G, Besada C, Badenes ML, Monforte AJ, Granell A. 2012. A non-targeted approach unravels the volatile network in peach fruit. PLoS One 7, e38992.

Seifikalhor M, Aliniaeifard S, Hassani B, Niknam V, Lastochkina O. 2019. Diverse role of γ-aminobutyric acid in dynamic plant cell responses. Plant Cell Reports 38, 847–867.

Shabala L, Zhang J, Pottosin I, Bose J, Zhu M, Fuglsang AT, Buendia AV, Massart A, Hill CB, Roessner U, Bacic A, Wu H, Azzarello E, Pandolfi C, Zhou M, Poschenrieder C, Mancuso S, Shabala S. 2016. Cell-type-specific H^+^-ATPase activity in root tissues enables K^+^ retention and mediates acclimation of barley (*Hordeum vulgare*) to salinity stress. Plant Physiology 172, 2445–2458.

Shahid SA, Zaman M, Heng L. 2018. Soil salinity: Historical perspectives and a world overview of the problem. In guideline for salinity assessment, mitigation and adaptation using nuclear and related techniques, Zaman M., Shahid, S. A., Heng, L. eds. (Cham, Switzerland: Springer), pp. 43–53.

Shahzad AN, Pitann B, Ali H, Qayyum MF, Fatima A, Bakhat HF. 2015. Maize genotypes differing in salt resistance vary in jasmonic acid accumulation during the first phase of salt stress. Journal of Agronomy and Crop Science 201, 443–451.

Shahzad R, Waqas M, Khan AL, Hamayun M, Kang SM, Lee IJ. 2015. Foliar application of methyl jasmonate induced physio-hormonal changes in *Pisum sativum* under diverse temperature regimes. Plant Physiology Biochemistry 96, 406–416.

Shaki F, Maboud HE, Niknam V. 2019. Effects of salicylic acid on hormonal cross talk, fatty acids profile, and ions homeostasis from salt-stressed safflower. Journal of Plant Interactions 14, 340–346.

Sharma A, Kumar V, Yuan H, Kanwar MK, Bhardwaj R, Thukral AK, Zheng B. 2018. Jasmonic acid observed treatment stimulates insecticide detoxification in *Brassica juncea* L. Frontiers in Plant Science 9, 1609.

Shi L, Guo MM, Ye NH, Liu YG, Liu R, Xia YJ, Cui SX, Zhang JH. 2015. Reduced ABA accumulation in the root system is caused by ABA exudation in upland rice (*Oryza sativa* L. var. Gaoshan 1) and this enhanced drought adaptation. Plant Cell Physiology 56, 951–964.

Shu S, Yuan Y, Chen J, Sun J, Zhang W, Tang Y, Zhong M, Guo S. 2015. The role of putrescine in the regulation of proteins and fatty acids of thylakoid membranes under salt stress. Scientific Reports 5, 14390.

Siddiqi KS, Husen A. 2019. Plant response to jasmonates: current developments and their role in changing environment. Bulletin of the National Research Centre 43, 153.

Singha KT, Sreeharsha RV, Marriboina S, Attipalli RR. 2019. Dynamics of metabolites and key regulatory proteins in the developing seeds of *Pongamia pinnata*, a potential biofuel tree species. Industrial Crops and Products 140, 111621.

Skorupa M, Gołębiewski M, Kurnik K, Niedojadlo J, Kesy J, Klamkowski K, Wójcik K, Treder W, Tretyn A, Tyburski J. 2019. Salt stress vs. salt shock the case of carbohydrate beet and its halophytic ancestor. BMC Plant Biology 19, 57.

Slama I, Abdell C, Bouchereau A, Flowers T, Savoure A. 2015. Diversity, distribution and roles of osmoprotective compounds accumulated in halophytes under abiotic stress. Annals of Botany 115, 433–447.

Spiess GM, Hausman A, Yu P, Cohen JD, Rampey RA, Zolman BK. 2014. Auxin input pathway disruptions are mitigated by changes in auxin biosynthetic gene expression in *Arabidopsis*. Plant Physiology 165, 1092–1104.

Sreeharsha RV, Shalini M, Singha KT, Attipalli RR. 2016. Unravelling molecular mechanisms from floral initiation to lipid biosynthesis in a promising biofuel tree species, *Pongamia pinnata* using transcriptome analysis. Scientific Reports 6, 34315.

Tavallali V, Karimi S. 2019. Methyl jasmonate enhances salt tolerance of almond rootstocks by regulating endogenous phytohormones, antioxidant activity and gas exchange. Journal of Plant Physiology 234-235, 98–105.

Thor K. 2019. Calcium-nutrient and messenger. Frontiers in Plant Science 10, 440.

Tognetti VB, Aken OV, Morreel K, Vandenbroucke K, van-de-Cotte B, De-Clercq I, Chiwocha S, Fenske R, Prinsen E, Boerjan W, Genty B, Stubbs KA, Inzé D, Breusegem FV. 2010. Perturbation of indole-3-butyric acid homeostasis by the UDP-glucosyltransferase *UGT74E2* modulates *Arabidopsis* architecture and water stress tolerance. The Plant Cell 22, 2660–2679.

Tuteja N. 2007. Abscisic ccid and abiotic stress signaling. Plant Signaling Behavior 2, 135–138.

Uddin MR, Thwe AA, Kim YB, Park WT, Chae SC, Park SU. 2013. Effects of jasmonates on sorgoleone accumulation and expression of genes for sorgoleone biosynthesis in sorghum roots. Journal of Chemical Ecology 39, 712–722.

Ueda J, Kato J. 1982. Inhibition of cytokinin-induced plant growth by jasmonic acid and its methylester. Physiologia Plantarum 54, 249–252.

Wang F, Guo Z, Li H, Wang M, Onac E, Zhou J, Xia X, Shi K, Yu J, Zhou Y. 2016. Phytochrome A and B function antagonistically to regulate cold tolerance via abscisic acid-dependent jasmonate signaling. Plant Physiology 170, 459–471.

Wang J, Li S, Gong X, Xu J, Li M. 2020. Functions of jasmonic acid in plant regulation and response to abiotic stress. International Journal of Molecular Sciences 21, 1446.

Wang YF, Munemasa S, Nishimura N, Ren HM, Robert N, Han M, Puzorjova I, Kollist H, Lee S, Mori I, Schroeder JI. 2013. Identification of cyclic GMP-activated nonselective Ca^2+^ permeable cation channels and associated CNGC5 and CNGC6 genes in *Arabidopsis* guard cells. Plant Physiology 163, 578–590.

Wei P, Wang L, Liu A, Yu B, Lam HM. 2016. *GmCLC1* confers enhanced salt tolerance through regulating chloride accumulation in soybean. Frontiers in Plant Science 25, 1082.

Wu H. 2018. Plant salt tolerance and Na^+^ sensing and transport. The Crop Journal 6, 215–225.

Wu H, Shabala L, Azzarello E, Huang Y, Pandolfi C, Su N, Wu Q, Cai S, Bazihizina N, Wang L, Zhou M, Mancuso S, Chen Z, Shabala S. 2018. Na^+^ extrusion from the cytosol and tissue-specific Na^+^ sequestration in roots confer differential salt stress tolerance between durum and bread wheat. Journal of Experimental Botany 69, 3987–4001.

Wu X, He J, Chen J, Yang S, Zha D. 2014. Alleviation of exogenous 6-benzyladenine on two genotypes of eggplant (*Solanum melongena* Mill.) growth under salt stress. Protoplasma 251, 169–176.

Xie X, He Z, Chen N, Tang Z, Wang Q, Cai Y. 2019. The roles of environmental factors in regulation of oxidative stress in plant. BioMed Research International 2019, 9732325.

Xu L, Zhao H, Ruan W, Deng M, Wang F, Peng J, Luo J, Chen Z, Yi K. 2017. ABNORMAL INFLORESCENCE MERISTEM1 functions in salicylic acid biosynthesis to maintain proper reactive oxygen species levels for root meristem activity in rice. The Plant Cell 29, 560–574.

Yang CY, Liang YB, Qiu DW, Zeng HM, Yuan JJ, Yang XF. 2018. Lignin metabolism involves *Botrytis cinerea* BcG1-induced defense response in tomato. BMC Plant Biology 18, 103.

Yang T, Lv R, Li J, Lin H, Xi D. 2018. Phytochrome A and B negatively regulate salt stress tolerance of *Nicotiana tobacum* via ABA-jasmonic acid synergistic cross-talk. Plant Cell Physiology 59, 2381–2393.

Yang Y, Guo Y. 2018. Elucidating the molecular mechanisms mediating plant salt-stress responses. New Phytologist 217, 523–539.

Yang Y, Qi M, Mei C. 2004. Endogenous salicylic acid protects rice plants from oxidative damage caused by aging as well as biotic and abiotic stress. The Plant Journal 40, 909–919.

Yong HY, Zou Z, Kok EP, Kwan BH, Chow K, Nasu S, Nanzyo M, Kitashiba H, Nishio T. 2014. Comparative transcriptome analysis of leaves and roots in response to sudden increase in salinity in *Brassica napus* by RNA-seq. BioMed Research International 2014, 467395.

Zelm EV, Zhang Y, Testerink C. 2020. Salt tolerance mechanisms of plants. Annual Review of Plant Biology 71, 24.1-24.31.

Zhang M, Cao Y, Wang Z, Wang ZQ, Shi J, Liang X, Song W, Chen Q, Lai J, Jiang C. 2018. A retrotransposon in an HKT1 family sodium transporter causes variation of leaf Na^+^ exclusion and salt tolerance in maize. New Phytologist 217, 1161–1176.

Zhang Z, Mao C, Shi Z, Kou X. 2017. The amino acid metabolic and carbohydrate metabolic pathway play important roles during salt-stress response in tomato. Frontiers in Plant Science 8, 1231.

Zhao C, Zayed O, Zeng F, Liu C, Zhang L, Zhu P, Hsu CC, Tuncil YE, Tao WA, Carpita NC, Zhu JK. 2019. Arabinose biosynthesis is critical for salt stress tolerance in *Arabidopsis*. New Phytologist 224, 274–290.

Zhao Q, Tobimatsu Y, Zhou R, Pattathil S, Gallego-Giraldo L, Fu C, Jackson LA, Hahn MG, Kim H, Chen F, Ralph J, Dixon RA. 2013. Loss of function of cinnamyl alcohol dehydrogenase 1 leads to unconventional lignin and a temperature-sensitive growth defect in *Medicago truncatula*. Proceedings of the National Academy of Sciences of the United States of America 110, 13660–13665.

